# A 4D GelMA-Cad/Gelatin Hydrogel for High-Resolution DMD Bioprinting of Vascularized Soft Tissue Models

**DOI:** 10.64898/2026.10.01.756011

**Authors:** Jason X. Wang, Claire A. McClain, Evan Krystofiak, Holly E. David, Nicole T. Marguerite, W. Ryland Hoskins, Haley F. Sutphen, Jonathan A. Kropski, A. Scott McCall, Brian O’Grady

## Abstract

Digital micromirror device (DMD)-based bioprinting enables high-resolution fabrication of hydrogel-based tissue constructs, but print fidelity and hydrogel mechanics are typically coupled through polymer concentration, light exposure, and crosslinking density. This creates a materials constraint for vascular bioprinting, where constructs must preserve microscale architecture during fabrication while supporting multicellular remodeling, lumen formation, and long-term tissue organization. To meet these requirements, we developed a 4D GelMA-Cad/gelatin hydrogel with time-dependent mechanics that maintains print fidelity while providing a soft, cell-permissive matrix. GelMA-Cad provides a covalently photocrosslinked, bioactive network containing an N-cadherin-mimetic peptide, while uncrosslinked gelatin serves as a transient matrix-modifying component that diffuses from the hydrogel after fabrication, producing further softening without bulk degradation of the GelMA-Cad network. We characterized hydrogel chemistry with NMR, quantified time-dependent changes in stiffness, visualized pore morphology with cryo-SEM, and evaluated the platform in retinal and pulmonary constructs. The 4D hydrogel supports 3D-patterned accuracy, cell viability, inner blood-retinal barrier vasculature, and vascularized lung-alveolar tissue models. Single-cell RNA-sequencing of the pulmonary vascular–alveolar model demonstrated compartment-specific transcriptional responses, including enrichment of lung capillary-associated endothelial features and AT1-like epithelial states. Overall, this platform decouples DMD print fidelity from post-print matrix mechanics for vascularized soft tissue bioprinting and tissue modeling.

## 1. Introduction

Advances in 3D bioprinting have enabled increasingly precise control over the spatial organization of cells and extracellular matrix-mimetic materials ^[1–3]^. This progress has expanded the ability to fabricate tissue constructs with defined geometry ^[4]^, multicellular patterning ^[5,6]^, and microscale vascular-like architectures ^[7,8]^, enabling larger and more physiologically relevant engineered tissues ^[9]^. Projection-based approaches, including digital micromirror device (DMD)-based bioprinting, are particularly useful for this purpose because patterned light can rapidly photocrosslink hydrogel precursors with high spatial resolution ^[10,11]^. Substantial progress has been made in bioink engineering by tuning hydrogel composition ^[12–16]^, photoinitiator systems ^[17–19]^, light exposure conditions ^[11,20]^, and crosslinking chemistry ^[21,22]^ to improve print fidelity while maintaining cytocompatibility ^[23]^. These strategies have enabled the development of printable hydrogel systems that incorporate bioactive ligands ^[24]^, degradable crosslinkers ^[25]^, and tissue-specific mechanical properties ^[21,26,27]^. In many cases, degradable or cell-remodelable hydrogels have provided an effective route to improve matrix permissiveness after printing by allowing cells to modify their surrounding microenvironments ^[28–31]^.

Despite these advances, the variables used to improve print fidelity also define the mechanical properties of the final hydrogel. Increasing polymer concentration ^[32]^, light exposure dose, or crosslinking density ^[33,34]^ can improve feature retention, but these parameters can also produce denser, stiffer networks that reduce matrix permissiveness, cell viability, and cell-driven remodeling ^[14,29,35]^. Conversely, softer hydrogel formulations may better support cellular integration, matrix remodeling, lumen formation, and multicellular organization, but can lack the mechanical stability required to maintain high-resolution features during fabrication, media changes, and long-term culture ^[17,36,37]^. A bioink strategy that preserves high-resolution features during fabrication while providing a mechanically permissive matrix during culture would address a central limitation in DMD bioprinting.

Vascularized tissue models are especially affected by this bioink design constraint because vascular network formation requires both defined architecture and matrix permissiveness ^[38–41]^. DMD bioprinting can provide the spatial control needed to pattern vascular architectures ^[4,42–45]^, but endothelial organization, lumen formation, mural-cell association, and extracellular matrix deposition depend on a hydrogel environment that permits cell spreading and remodeling after fabrication ^[46,47]^. Stiffer or highly crosslinked matrices may preserve printed geometry, but can restrict the cellular reorganization, survival, and multicellular interactions required for vascular network formation ^[48]^, whereas softer matrices may support vascular remodeling ^[49]^ but compromise structural fidelity and construct handling. Thus, vascular bioprinting provides a stringent test of whether a hydrogel can preserve microscale architecture during fabrication while supporting multiple interacting cell types, vascular remodeling, and tissue organization during long-term culture.

Here, to decouple high-resolution DMD print fidelity from the cell-permissive mechanical properties required for tissue formation, we developed a temporally tunable 4D GelMA-Cad/gelatin hydrogel for bioprinting soft tissue constructs. In this system, the GelMA-Cad network provides a patternable, covalently photocrosslinked, bioactive scaffold containing an N-cadherin-mimetic peptide, while unmodified gelatin functions as a transient matrix-modifying component that diffuses from the hydrogel after fabrication. We chemically characterized the hydrogel components by ¹H NMR, incorporated Curcumin-Na as an established water-compatible photoabsorber for controlling light-mediated hydrogel patterning, and validated an indentation-based approach for measuring apparent Young’s modulus in bioprinted hydrogels. We then quantified time-dependent changes in hydrogel stiffness, evaluated DMD print fidelity using complex acellular geometries, and assessed long-term cell-mediated remodeling within patterned constructs. Finally, we applied the platform to bioprinted retinal vascular tricultures that form lumenized inner blood-retinal barrier (iBRB)-like networks and pulmonary vascular–alveolar constructs that develop lumenized vascular beds and AT1-like epithelial states, with scRNA-seq used to define compartment-specific transcriptional responses in the pulmonary model. Overall, this work establishes a 4D hydrogel strategy that maintains microscale print fidelity during fabrication while providing a soft, cell-permissive matrix that undergoes time-dependent mechanical and microstructural remodeling during culture.

## 2. Methods

### 2.1 Lung Cell Culture

Human Lung Microvascular Endothelial Cells (HULEC-5a; CRL-3244, ATCC) were cultured in MCDB 131 (10372-019, Gibco) supplemented with 5 ng/ml epidermal growth factor (100-15, PeproTech), 10 ug/ml Hydrocortisone (A16292.03, Thermo Scientific), 10% FBS (A56704-02, Gibco), and 50 ug/ml gentamicin (15710064, Gibco). Primary Lung Fibroblasts, Normal, Human (HLF; PCS-201-013, ATCC) were cultured in Fibroblast Basal Medium supplemented with a low-serum Fibroblast Growth Kit (PCS-201-041, ATCC). HULEC-5a and HLF were cultured on tissue culture flasks that were pre-coated with 0.2% w/v gelatin for at least 3 h prior to cell seeding. At 80% confluency, cells were passaged using TrypLE Express (12604021, Thermofisher) for subculturing. Human lung organoids were derived from declined donors per VUMC IRB #192004. Organoids were cultured in domed Matrigel (356231, Corning) droplets to confluence in PneumaCult Alveolar Organoid Expansion Media (100-0847, Stemcell Technologies). Organoids were passaged by adding 2 mg/ml Dispase II (D4693-1G, Sigma Aldrich) and incubating at 37 °C for 1 hour with gentle disruption after 30 minutes to liberate whole organoids from the Matrigel droplets. Organoids were then washed 3 times with high glucose DMEM (21063-029, Gibco) to remove Dispase II and reseeded in Matrigel droplets at a 1:1 ratio with PneumaCult Alveolar Organoid Expansion Media supplemented with 10 uM Y-27632 dihydrochloride (Y0503, Sigma Aldrich). All cultures were incubated at 37 °C and 5% CO₂, with media refreshes every 2-3 days.

### 2.2 Retinal Cell Culture

Primary human retinal microvascular endothelial cells (ACBRI 181, Cell Systems) and primary human retinal pericyte cells (ACBRI 183, Cell Systems) were cultured in Complete Classic Medium with Serum and CultureBoost (4Z0-500, Cell Systems). Normal human dermal fibroblasts (PCS-201-012, ATCC) were maintained in Fibroblast Basal Medium supplemented with a low-serum Fibroblast Growth Kit (PCS-201-041, ATCC). Tissue culture flasks were pre-coated with 0.2% w/v gelatin for at least 3 h prior to cell seeding. All cultures were incubated at 37 °C and 5% CO₂, with media refreshes every 2-3 days. At 80% confluency, cells were passaged using TrypLE Express (12604021, Thermofisher) for subculturing. For the fabrication of 3D bioprinted constructs, the three cell lines were pooled into a triculture suspension within the bioink at a 4:1:1 ratio (endothelial:pericyte:fibroblast) to yield a total cellular density of 6 × 10⁶ cells/mL. Post-fabrication, the 3D bioprinted retinal constructs were maintained in Complete Classic Medium with daily media changes.

### 2.3 3D Multicellular Media

A custom culture medium, termed 3D multicellular media (3DMM), was derived from O’Grady et al, 2026 ^[50]^. In brief, the media was formulated using human endothelial serum-free media (hESFM; 11111044, ThermoFisher) as the basal medium and was supplemented with 250 ng/mL heat-stable basic fibroblast growth factor (bFGF; PHG0367, Gibco), 250 ng/mL vascular endothelial growth factor (VEGF; 100-20, PeproTech), 500 ng/mL epidermal growth factor (EGF; 100-15, PeproTech), 500 ng/mL of BPC-157/TB-4 peptide blend (SimplePeptide), 25 µg/mL fibronectin (33016015, ThermoFisher), and 50 µg/mL gentamicin (15710064, Gibco). 3DMM media was made fresh before use at a final volume of 20 mL.

### 2.4 Gelatin preparation

A 10% w/v gelatin stock solution was prepared by dissolving Type A gelatin (300 Bloom; G2500, Sigma-Aldrich) in 200 mL of DPBS (21-031-CV, Corning). The gelatin powder was gradually introduced to DPBS pre-heated to 50 °C under continuous magnetic stirring (500 rpm) and maintained for 2 hours to ensure complete dissolution. The warm solution was sterilized via vacuum filtration using a 0.22 µm polyethersulfone (PES) Nalgene Rapid-Flow unit (09-741-01, Thermofisher), aliquoted into 10 mL volumes, and stored at 4 °C for up to 6 months.

### 2.5 GelMA synthesis

Gelatin methacryloyl (GelMA) was synthesized following an established protocol ^[51]^. Briefly, Type B gelatin (250 Bloom; G9391, Sigma-Aldrich) was dissolved at 10% w/v in a 0.25 M carbonate-bicarbonate buffer (pH 9.4) at 55 °C. Methacrylic anhydride (276685, Sigma-Aldrich) was added dropwise, and the methacrylation reaction proceeded for 1 hour at 55 °C under constant stirring. The reaction was terminated by neutralizing the solution pH to 7.4 with 1 M HCl. The crude mixture was vacuum filtered through a pre-warmed 0.2 µm polyethersulfone (PES) membrane. The mixture was dialyzed using tangential flow filtration (TFF) using a Masterflex L/S (13-200-007, Fisher Scientific) with a 10K Minimate Omega membrane (50-197-9560, Fisher Scientific) against Milli-Q water at 50 °C for 16 hours to remove unreacted constituents. The purified GelMA solution was subsequently frozen stepwise to promote large crystal formation. The frozen solution was lyophilized at -80 °C and <0.1 mbar (75999-788, Labconco) for 5 days, and cryomilled via a liquid nitrogen-chilled polymer grinder (Z341789, Sigma) into a fine powder for long-term storage at -80 °C.

### 2.6 Curcumin-Na synthesis

Curcumin sodium (Curcumin-Na) was synthesized following a modified procedure adapted from He et al, 2023 ^[52]^. Briefly, 368 mg of curcumin (B21573-09, ThermoFisher) was dissolved in 25 mL of ethanol (DSP-MD.43, Decon Lab.). Subsequently, 15 mL of Milli-Q (MQ) water (18.2 MΩ·cm) and 294 mg of sodium bicarbonate (S6014, Sigma) were added, and the mixture was stirred at 300 rpm for 2 hours at room temperature. The solvent was evaporated overnight on a 37°C warming plate in a fume hood. The resulting dried Curcumin-Na was resuspended in 40 mL of MQ water and passed through a sterile 0.2 µm polyethersulfone (PES) syringe filter to remove undissolved particulates. The purified filtrate was aliquoted (≤15 mL per tube) and subjected to a stepwise freezing protocol to promote large ice crystal formation: 4 °C for ≥3 hours, -20 °C for ≥2 hours, and -80 °C for ≥4 hours. The samples were then lyophilized at -80 °C and <0.1 mbar (75999-788, Labconco) for 3 days and stored at -80 °C until use.

### 2.7 Synthesis of Methacryloyl-Functionalized N-Cadherin Peptide

A custom methacryloyl-functionalized N-cadherin-mimetic peptide was synthesized by Biomatik and GenScript for covalent incorporation into GelMA-based hydrogels. The peptide contained the N-cadherin HAVDI adhesion motif flanked by four glycine spacers on each side, with the core sequence K[Meth]GGGGHAVDIGGGGK[Meth]. Methacryloyl-L-Lysine (K[Meth]) residues were incorporated at both terminal positions, generating a dual-methacryloyl peptide with polymerizable handles at the N- and C-terminal ends. These methacryloyl groups enabled covalent incorporation of the peptide into the GelMA network during light-mediated photocrosslinking, thereby generating GelMA-Cad hydrogels. Peptides were synthesized at >98% purity and supplied as lyophilized 5 mg aliquots. For experimental hydrogel formulations, peptides were dissolved directly into the GelMA precursor solution prior to bioprinting at a final concentration of 5 mg/mL.

### 2.8 GelMA/gel N-cad 4D Hydrogel Formulation

Bioink formulations were prepared to achieve final macromer concentrations of 3.75% w/v gelatin methacryloyl (GelMA) and 2.5% w/v gelatin. Briefly, GelMA and the photoinitiator lithium phenyl-2,4,6-trimethylbenzoylphosphinate (LAP, 1% w/v final concentration for all acellular and retinal cultures while a 0.5% w/v final concentration was used for the pulmonary cultures) were dissolved in either cell culture media (for cellular prints) or Dulbecco’s Phosphate-Buffered Saline (DPBS, for acellular prints) at 37 °C under continuous agitation. Pre-warmed gelatin stock was then incorporated into the mixture, followed by the addition of Curcumin-Na to achieve a final concentration of approximately 1 mM (as determined via absorbance quantification). The resulting composite solution was sterilized via pre-warmed 0.2 µm syringe filtration. For cellular bioinks, synthetic N-Cadherin peptide was mixed into the filtered solution at a final concentration of 5 mg/mL, and cells were subsequently suspended in the bioink. For acellular control prints, the bioink was supplemented with an equivalent volume of DPBS in place of the cell and peptide suspensions to ensure the final macromer concentrations remained constant.

### 2.9 Print Probe Fabrication

Bioprinter probes were fabricated on a Formlabs Form 4 3D printer using Formlabs Black Resin V5 at a print resolution of 0.05mm z-height. Following printing, the probes were washed in an automated isopropyl alcohol (IPA) bath for 20 minutes, rinsed with fresh IPA, and air-dried for 10 minutes. After support structure removal, the printed components were post-cured in a UV chamber for a minimum of 30 minutes at 60 °C. Glass slides (260379-12, Ted Pella) were cleaned with ethanol and Milli-Q (MQ) water, dried, and adhered to the probe rims using a thin layer of Loctite SI 5056 adhesive (1214249, Henkel Adhesives) applied via a disposable micro applicator brush. The adhesive was set using a Thorlabs 385nm UV curing system (CS20K3, Thorlabs) for 10 minutes at 337 mW/cm^2^ and subsequently cured at room temperature for 3 days. Excess glue was gently scraped off using a razor blade. The assembled probes were washed sequentially with IPA and MQ water prior to storage at room temperature.

### 2.10 Probe Silanization

The glass surfaces of 3D-printed probes were sequentially cleaned with acetone and Milli-Q (MQ) water and air dried. A silanization solution was prepared by adjusting a solvent mixture of 95% ethanol and 5% deionized water (DIW) to a pH of 4.0–5.0 using glacial acetic acid, followed by the addition of 2% v/v hexyltrimethoxysilane (HTMS) (A428690, AmBeed). This solution was agitated on an orbital shaker at 100 rpm for 10 minutes to facilitate HTMS hydrolyzation. The probes were plasma-treated for 10 minutes, then immediately submerged in 4 mL of the hydrolyzed silane solution within a 6-well plate and agitated at 100 rpm for 5 minutes on an orbital shaker. After incubation, the probes were transferred to an oven and baked at 110–120 °C for 20 minutes. Post-baking, the probes were washed three times with anhydrous ethanol, rinsed with MQ water, and air-dried. Hydrophobic coating efficacy was evaluated utilizing a friction angle tester by depositing a 20 µL droplet of MQ water onto the probe surface; probes demonstrating initial droplet movement at tilt angles <45° were considered successfully functionalized and retained for printing.

### 2.11 Bioprinting

Bioprinting was performed using a Cellink Bionova X bioprinter. Prior to printing, the chamber, methacrylated well plates (D16110025296, CELLINK), and custom hydrophobic printing probes were sterilized via 5 minutes of UVC irradiation (270 nm). All required reagents and hardware were pre-warmed to 37 °C. To prevent gel adhesion, probes were surface-treated with 200 µL of sterile-filtered 5% w/v Pluronic F-127 solution (P2443, Sigma-Aldrich) for 1 minute, after which the excess solution was carefully pipetted off. Hydrogel solution was loaded into well plates under sterile conditions. Digital models were sliced at a 10 µm layer thickness. Photocrosslinking was performed using 405 nm UV light at 16 mW/cm² (100% intensity) with a 5-second exposure time per layer. Printer movement settings were configured with a 4-second rest time post-lowering, a 0.5 mm/s lowering speed, a 0.5 mm/s retract speed, and a constant 2.0 mm lowering and retract distance. Following fabrication, uncrosslinked bioink was recovered, and the constructs were washed with 1 mL of warmed DPBS (21-031-CV, Corning) to remove residual gel and unattached cells before adding culture media.

### 2.12 Mechanical Indentation

Mechanical testing via ball bearing indentation was adapted from established methodologies ^[53,54]^. Bioprinted hydrogel discs (6.5 mm diameter, 2.0 mm height) were maintained in Dulbecco’s Phosphate-Buffered Saline (DPBS; 21-031-CV, Corning) with 0.1% w/v sodium azide (014314-09, ThermoFisher). To visualize deformation, an aqueous 1.0 µm red fluorescent particle suspension (Ex/Em 542/612 nm; R0100, ThermoFisher) was applied to the construct surface. A 1 mm diameter Aluminum 2017 ball bearing (34665k35, McMaster-Carr) was carefully centered on the hydrogel, and additional fluorescent particles were deposited directly onto the bearing. Indentation depth was optically quantified using a Zeiss LSM 980 confocal microscope equipped with a 37 °C humidified environmental chamber. Z-stack imaging (Alexa Fluor 555, 10 µm pinhole) was used to measure the vertical distance from the bearing’s lowest focal point to the undeformed hydrogel surface. The gels were kept in solution with only the top surface exposed to air during the measurement to avoid any complications with mechanical stiffening due to dehydration while also avoiding bearing buoyancy and water surface tensions.

### 2.13 Nuclear Magnetic Resonance Analysis

^1^H-NMR spectra were acquired using a 400 MHz Bruker Avance Neo spectrometer. Aqueous samples were prepared by dissolving 10 mg of analyte in 580 µL of deuterium oxide (D_2_O; 151882, Sigma-Aldrich) supplemented with 20 µL of a TMSP internal standard solution (3 mg TMSP [269913, Sigma-Aldrich] in 600 µL D_2_O). Non-aqueous samples were prepared by dissolving 10 mg of analyte in 600 µL of deuterated dimethyl sulfoxide (DMSO-d6; 570672, Sigma-Aldrich). Solutions were centrifuged to eliminate air bubbles and transferred to glass NMR tubes (WG-1000-7, Wilmad-LabGlass) for analysis.

### 2.14 Live & Dead Assay

Bioprinted cellular constructs were stained with calcein AM (C34852, Thermofisher) for live cells and propidium iodide (P3566, Thermofisher) for dead cells, both at a concentration of 10 uM. Cells were stained for 15 minutes while placed on a rocker (88-861-025, Fisher Scientific) in the incubator at 37 °C and 5% CO_2_. Cells were then washed with DPBS for 10 minutes. Cell constructs were then confocal imaged in an environmentally controlled chamber with 100% humidity, 37 °C, and 5% CO_2_ (Zeiss LSM 980).

### 2.15 Immunofluorescence staining

Bioprinted hydrogels were fixed in 4% paraformaldehyde in Dulbecco’s Phosphate-Buffered Saline (DPBS; 21-031-CV, Corning) for 30 s and washed three times in DPBS for 15 min each. Samples were blocked and permeabilized overnight at 4 °C in a buffer containing 5% goat serum (50062Z, ThermoFisher), 0.01% v/v Triton X-100 (A16046-AE, Thermofisher), and 0.1% w/v sodium azide (014314-09, Thermofisher). Constructs were then incubated overnight at 4 °C in a 1 mL mixture of pre-conjugated antibodies and fluorescent probes: 2.5 µg/mL PDGFRβ (53-1402-82, Invitrogen), 2.0 µg/mL VE-cadherin (CL488-83766-2, ProteinTech), 1.0 µg/mL Collagen IV (51-9871-82, Invitrogen), 0.5 µg/mL phalloidin (A30105, Invitrogen), 2.0 µg/mL Lectin-LEA (L32470, Invitrogen), 1.0 µg/mL EpCAM (324228, Biolegned), 2.0 µg/mL PODXL (CL48818150, ProteinTech) and 1.0 µg/mL DAPI (62248, ThermoFisher). Following staining, the hydrogels were washed twice in DPBS for 15 min each prior to imaging.

### 2.16 Cryo-Scanning Electron Microscopy

For freeze-etching, hydrogels were vitrified using a Leica EM-ICE high-pressure freezer. Hydrogels were placed in 3.0 mm domed planchettes (Leica #16770132). All remaining space in the planchettes was filled with 2-methylpentane as filler. After vitrification, the samples were loaded onto a Leica ACE600 freeze fracture device using a Leica VCM/VCT vacuum transfer station. Samples were brought to -110 °C and fractured using a cold knife, after fracturing the knife cold knife was placed directly over the fractured surface while the samples were brought to -100 °C for 20 minutes to remove the 2-methylpentane and etch away surface water. After Etching the samples were shadowed with 3 nm of ebeam deposited platinum at 45° relative to the sample and 5 nm carbon at 90°. Samples were cryo-transferred to a Zeiss Crossbeam 550 equipped with a Leica VCT cryostage. Samples were imaged at 2 keV at 50 nA using the lower secondary electron detector using drift corrected frame averaging.

### 2.17 scRNA-seq Preparation

Cells were liberated from the 4D hydrogel by incubating 2 mg/ml Dispase II (D4693-1G, Sigma Aldrich) at 37 °C for 10 mins. The cell mixture was then washed three times with DMEM (21063-029, Gibco) and the Animal Component-Free Cell Dissociation Kit (5426, Stemcell Technologies) was used to singularize the organoids to create a single cell suspension. The sample was then washed once with DPBS and prepared for scRNA seq using the GEM-X Flex Sample Preparation v2 Kit (1000781, 10x Genomics) following 10x Genomics Demonstrated Protocol CG000782 | Rev D Fixation of Cells & Nuclei for GEM-X Flex Gene Expression.

### 2.18 scRNA-seq Analysis

Processing of the demultiplexed FASTQs was performed using ScanPy and Rapids Single Cell and standard workflows. Briefly, initial QC included 200-8000 genes expressed and <10% mitochondrial count content. Doublets were removed using Scrublet. Highly variable genes were used for PCA analysis and initial clustering. Cell compartments were separated, reclustered,and differential gene expression performed by compartment by rank_gene_groups usings Wilcoxon testing. Composite scores were calculated using score_genes and gene sets derived from cited references.

## 3. Results & Discussion

### 3.1 Engineering Time-Dependent Softening GelMA-Cad/Gelatin Hydrogels for DMD Bioprinting

The central objective of this study is to develop a bioink platform that resolves a key materials challenge in DMD bioprinting: preserving mechanical robustness during patterned-light fabrication while transitioning after printing into a softer, cell-permissive microenvironment capable of supporting multicellular remodeling and vascular network formation. The overall strategy is summarized in **Figure 1A**, where bioink precursors and cells are combined for in-well DMD bioprinting, cultured to generate multicellular vascularized tissue constructs, and subsequently characterized by fluorescence imaging and single-cell RNA sequencing. To enable this workflow, we engineered a temporally tunable 4D GelMA-Cad/gelatin hydrogel in which the structural, bioinstructive, and transient matrix-modifying functions of the material were distributed across distinct polymer components (**Figure 1B**).

**Figure 1.**
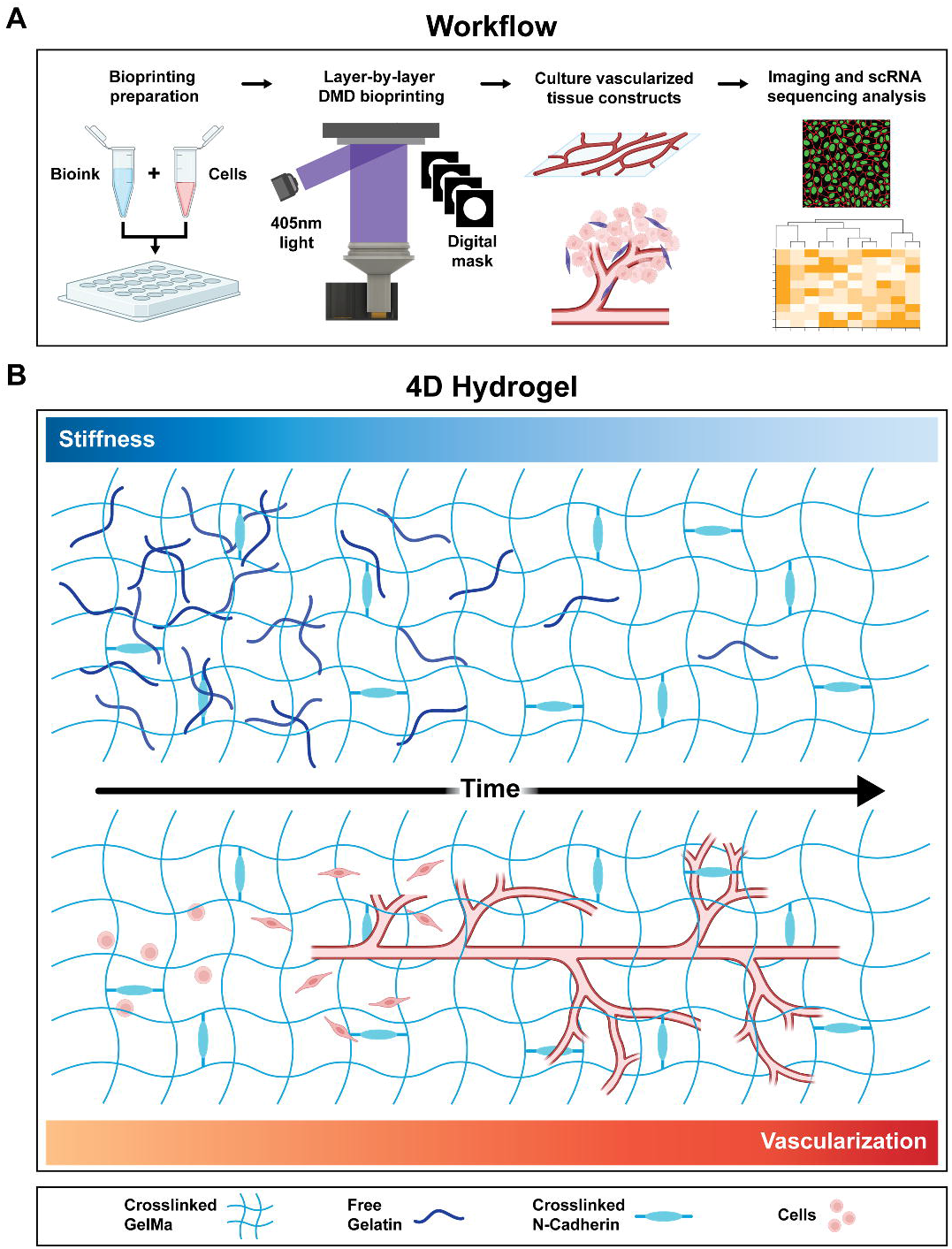
DMD-bioprinted 4D GelMA-Cad/gelatin hydrogel supports vascularization of tissue-mimetic constructs. A) Schematic overview of the in-well DMD bioprinting workflow. Bioink precursors and cells are combined directly within a 24-well plate for layer-by-layer DMD bioprinting, generating multicellular vascularized tissue constructs that are subsequently cultured and analyzed by fluorescence imaging and single-cell RNA sequencing. B) Schematic overview of the 4D GelMA-Cad/gelatin hydrogel and its time-dependent transition from print-supportive stiffness to vascularization-permissive remodeling. A photocrosslinked GelMA-Cad network containing covalently incorporated N-cadherin-mimetic peptides provides a stable bioactive scaffold, while unmodified gelatin is physically entrapped as a transient, matrix-modifying component. Initially, the entrapped gelatin increases the effective matrix stiffness, enabling high-fidelity microfeature patterning during DMD bioprinting. Over time, gelatin diffuses from the hydrogel, producing a temporal reduction in matrix stiffness while preserving the initially patterned architecture. This time-dependent matrix remodeling generates a more cell-permissive environment that supports cell viability, vascular remodeling, and time-dependent vascular network formation within the bioprinted constructs.

GelMA provides the stable photocrosslinked scaffold, while the photocrosslinkable N-cadherin-derived peptide is incorporated to introduce bioinstructive cell–cell adhesion cues that support multicellular organization and vascular network formation ^[50,55,56]^. In parallel, unmodified gelatin functions as a physically entrapped, transient matrix-modifying component that increases the initial effective stiffness of the overall hydrogel during fabrication and then gradually diffuses from the matrix during culture. This coordinated temporal transition allows the printed construct to begin in a print-supportive state, where increased stiffness promotes structural fidelity, before shifting toward a softer and more cell-permissive matrix as gelatin disseminates from the scaffold. As the matrix becomes more permissive, vascular organization increases over time, linking the material’s time-dependent mechanical remodeling to the biological remodeling required for vascular network formation. This material concept was then translated into the in-well DMD bioprinting workflow shown in **Figure 2A**, where a homogeneous cell-laden precursor solution is prepared directly for patterned-light fabrication using defined printer settings and digital masks containing positive and negative microfeatures.

**Figure 2.**
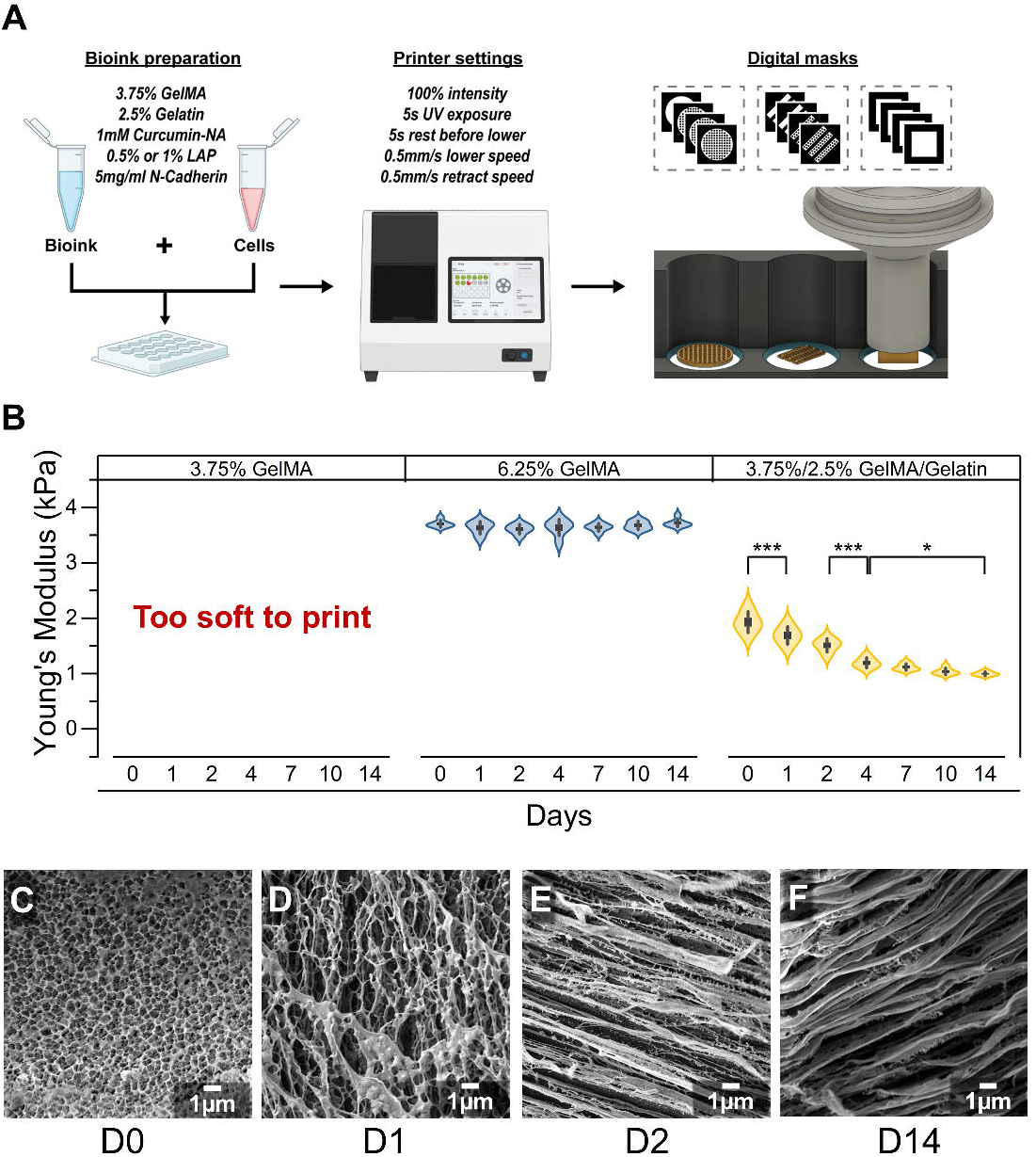
GelMA-Gelatin Hydrogel Bioprinting Workflow and Characterization of 4D Properties. A) Workflow for DMD bioprinting of patterned hydrogel constructs. Bioink was prepared using the described formulation, combined with cells, and transferred into a 24-well plate for in-well bioprinting. DMD bioprinter parameters were then defined for subsequent fabrication experiments. Digital masks were used to guide layer-by-layer bioprinting of specified geometries containing both positive and negative microfeatures. B) Apparent Young’s modulus of acellular bioprinted hydrogel formulations quantified via aluminum ball bearing macroindentation over a 14-day culture period. The 3.75% w/v GelMA formulation lacked sufficient mechanical stability for bioprinting. The 6.25% w/v GelMA control maintained a consistent stiffness (∼3.66 kPa, +/-0.095) across the time course. In contrast, the 4D hydrogel (3.75% w/v GelMA / 2.5% w/v gelatin) exhibited significant temporal softening, relaxing from an initial stiffness of 1.93 ± 0.19 kPa at Day 0 to .997 ± 0.036 kPa by Day 14. (C–F) Representative cryo-scanning electron microscopy (cryo-SEM) images revealing the microstructural progressive pore opening of the 4D hydrogel immediately after printing at (C) Day 0, and subsequently at (D) Day 1, (E) Day 2, and (F) Day 14. (Scale bars = 1 µm). Data in (B) are presented as violin plots showing the raw data distribution, with internal box plots representing the standard error (SE) and whiskers representing the standard deviation (SD) (n = 6 independent samples per formulation). Statistical analysis was done by two-way ANOVA testing, where *P<0.05, **P<0.01, ***P<0.001, and P>0.05 (no significant difference). See supplementary Table 1 for Tukey pairwise comparison groupings and descriptive statistics.

To translate this material design into a reproducible DMD bioink, the precursor formulation was optimized around three integrated components: the GelMA-Cad/gelatin matrix, LAP-mediated radical polymerization, and Curcumin-Na as a water-compatible photoabsorber (**Figure 2A**). This formulation strategy is critical because high-resolution DMD bioprinting depends on both the projected digital mask and the spatial confinement of radical-mediated crosslinking within the precursor solution. Because LAP is distributed throughout the bioink, radicals generated during UV exposure can diffuse or propagate beyond the illuminated region and initiate GelMA crosslinking outside the intended patterned geometry, reducing feature definition around small negative spaces. Curcumin-Na was therefore incorporated as an established optical-control strategy to attenuate off-target radical-mediated polymerization while maintaining an aqueous, cell-compatible formulation ^[52]^. Conversion of native curcumin to the sodium salt enabled homogeneous incorporation into the precursor without organic co-solvents, and because Curcumin-Na remained unbound after photocrosslinking, the compound washed out during culture, leaving the printed construct defined primarily by the GelMA-Cad scaffold and transient gelatin-modified matrix. The chemistry required for this formulation was confirmed by ¹H NMR, with GelMA synthesis supported by methacryloyl-associated proton peaks at ∼5.4–5.8 ppm and methyl proton peaks at ∼1.8–2.2 ppm, consistent with successful methacryloyl substitution of gelatin **(Figure S1)**. Curcumin-Na synthesis was similarly supported by loss of the phenolic hydroxyl resonance near ∼9.65 ppm and shifts associated with the keto-enol/curcumin backbone near ∼6.1 ppm, consistent with conversion of curcumin into the water-compatible sodium salt form **(Figure S2)**.

With the formulation chemistry established, we quantified the time-dependent mechanical behavior of printed constructs using aluminum ball-bearing microindentation and confocal Z-stack imaging. This method enabled optical measurement of indentation depth in soft, hydrogel discs without requiring bulk compression or extensive sample handling. The deformation profile was analyzed using a modified Hertz model to calculate apparent Young’s modulus, with values reported as apparent moduli due to the model assuming a Poisson’s ratio for the hydrogel network **(Figure S3)**. Visual and topographical validation further confirmed that the ball-bearing setup produced a measurable, spatially defined indentation profile suitable for comparing formulation-dependent and time-dependent stiffness changes **(Figure S4)**. Using this approach, we compared 3.75% w/v GelMA-Cad, 6.25% w/v GelMA-Cad, and the 4D formulation composed of 3.75% w/v GelMA-Cad with 2.5% w/v unmodified gelatin. This comparison separated the effect of total polymer content from the effect of gelatin as a transient matrix-modifying component. The 3.75% GelMA-Cad formulation was too soft to print reliably and failed to maintain the target 2mm construct height, while the 6.25% GelMA-Cad control printed successfully, but remained mechanically static at 3.66 ± 0.095 kPa across 14 days (**Figure 2B**). In contrast, the 4D GelMA-Cad/gelatin hydrogel printed as an intact construct, then underwent significant time-dependent softening, decreasing from 1.93 ± 0.19 kPa at Day 0 to 0.997 ± 0.036 kPa by Day 14 (**Figure 2B; Table S1)**. This behavior supports the intended role of gelatin as a transient mechanical modifier rather than a permanent stiffening phase. The added gelatin increased the effective stiffness needed for print-stage handling and construct formation, while its gradual diffusion from the scaffold allowed the matrix to transition after fabrication toward a softer, more cell-permissive state. This transient reinforcement produced the intended 4D mechanical profile: an initially stable printed construct capable of maintaining geometry during fabrication, followed by progressive softening toward a matrix state expected to support pore opening and subsequent cell-driven remodeling.

The time-dependent mechanical behavior of the 4D hydrogel was paralleled by changes in matrix microstructure, as shown by cryo-SEM across the same culture period (**Figure 2C–F**). Immediately after printing, the hydrogel exhibited a dense polymer network, consistent with a matrix capable of maintaining construct integrity during DMD fabrication (**Figure 2C**). By Day 1, larger void spaces appeared throughout the scaffold as the physically entrapped gelatin began to disengage from the covalently crosslinked GelMA-Cad network (**Figure 2D**). This microstructural transition became more pronounced by Day 2, where larger channel-like features were distributed through the hydrogel, supporting the interpretation that gelatin elution increased internal matrix space while preserving the primary photocrosslinked scaffold (**Figure 2E**). The open architecture continued through Day 14, indicating that the reduction in apparent modulus measured mechanically corresponded to a stable matrix transition rather than loss of construct integrity (**Figure 2F**). These structural changes support the proposed 4D material mechanism: unmodified gelatin contributes to the initial post-print matrix state and then gradually exits the scaffold, generating a softer and more open GelMA-Cad network. By increasing matrix permissiveness through a transient gelatin phase rather than degradation of the primary photocrosslinked scaffold, this strategy preserves construct stability while creating a microenvironment more favorable for cellular remodeling, matrix integration, and vascular network formation.

These material characterization studies establish the GelMA-Cad/gelatin hydrogel as a temporally tunable and bioinstructive bioink for DMD bioprinting. The formulation integrates a chemically validated photocrosslinkable GelMA-Cad scaffold, covalently incorporated N-cadherin-derived adhesion cues, a transient gelatin phase, Curcumin-Na-mediated optical and radical control, and validated time-dependent mechanical softening. This combination addresses the competing requirements of patterned-light fabrication and vascularized tissue culture by providing construct-level stability during printing, followed by transition into a softer and more open matrix state that supports post-print cellular remodeling. This material foundation then enabled evaluation of whether the same 4D hydrogel strategy could preserve complex printed geometries while supporting long-term cell-mediated remodeling within patterned constructs.

### 3.2 The 4D Hydrogel Supports High-Fidelity DMD Patterning and Cell-Mediated Matrix Remodeling

Having established that the GelMA-Cad/gelatin hydrogel undergoes time-dependent softening without loss of construct integrity, we evaluated whether the same formulation could preserve the spatial fidelity encoded by DMD projections during fabrication. This distinction is critical because bulk mechanical characterization alone does not define printability; DMD bioink performance also requires accurate reproduction of microscale positive and negative features while resisting deformation during repeated exposure, immersion, and probe movement. Three CAD geometries were therefore selected to challenge distinct failure modes in the printing process (**Figure 3A–C; Figure S5**). The bicycle geometry assessed local resolution and features bridging within closely spaced spoke patterns, where radical-mediated overcuring would be most evident. The “HI” geometry assessed dimensional fidelity across 100, 150, and 200 µm beam widths while simultaneously challenging aspect-ratio stability at a uniform print height. Finally, the honeycomb lattice evaluated construct-level robustness in a larger continuous-wall architecture exposed to cupping, suction, and blowout forces during layer-by-layer DMD fabrication. All three acellular geometries were fabricated using the optimized 4D GelMA-Cad/gelatin formulation to a target height of 200 µm, with 10 µm layers, 5 s layer exposures, and 16 mW/cm² light intensity at 405 nm (**Figure 3D–F**). Constructs were analyzed 24 h after printing using intrinsic hydrogel autofluorescence and confocal microscopy, allowing geometric fidelity to be evaluated after removal of uncrosslinked precursor, washout of soluble components, and early swelling equilibration. This timing provided a more stringent assessment than immediate post-print imaging, because the constructs had already undergone the initial post-fabrication handling steps that can reveal weak feature retention or mechanical instability.

**Figure 3.**
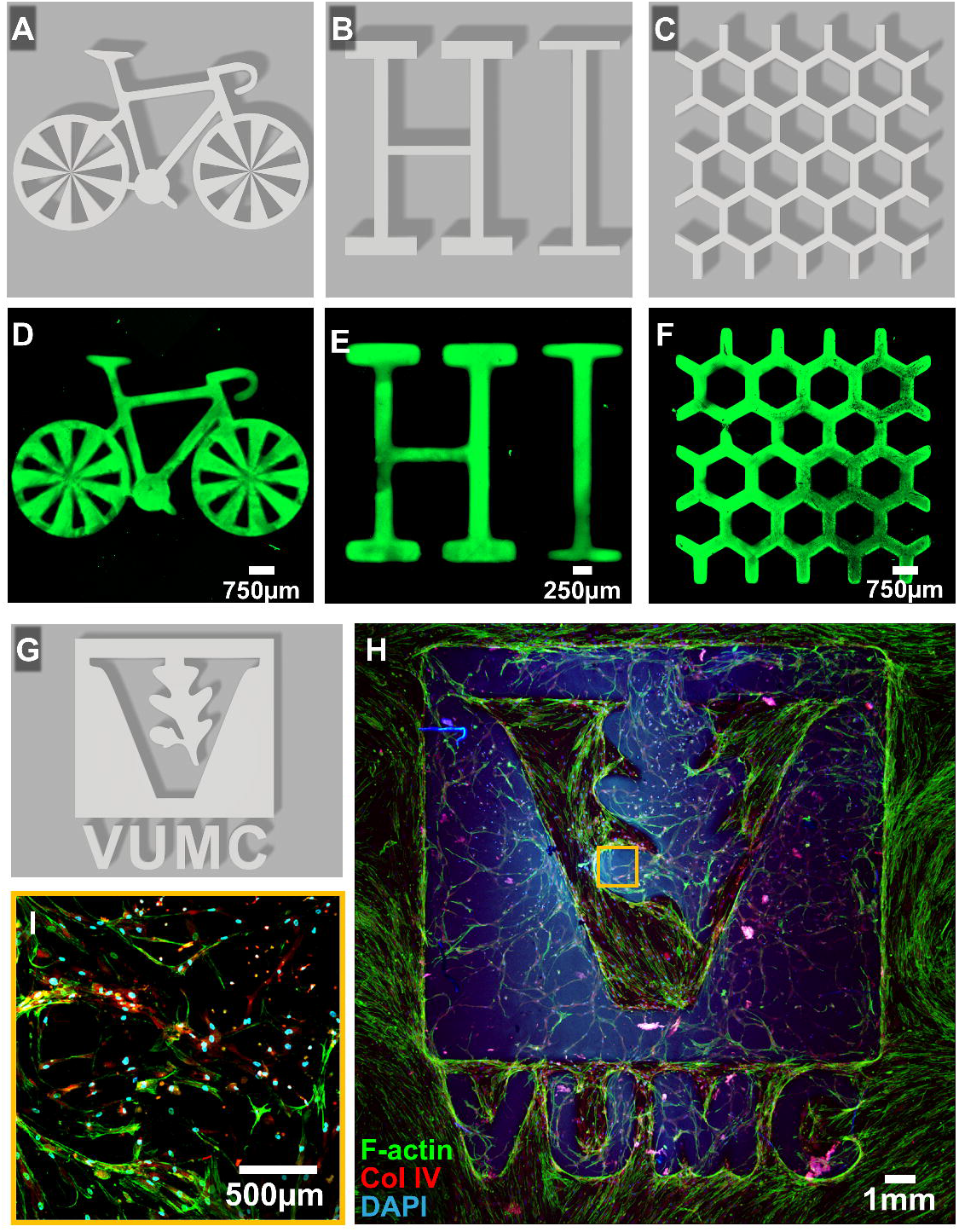
High-fidelity digital micromirror device (DMD) bioprinting and cellular remodeling of the 4D hydrogel. (A–C) Computer-aided design (CAD) renderings of complex geometries designed to evaluate specific optical and mechanical challenges during DMD bioprinting: (A) a bicycle model to assess the print resolution of directly adjacent positive and negative features; (B) a “HI” construct with variable dimensions to test structural stability against column effects and assess resulting swelling ratios; and (C) a honeycomb lattice to evaluate macroscopic resilience against vertical cupping and blowout forces present across larger patterns. (D–F) Maximal Z-projections of corresponding acellular 4D hydrogel bioprints visualized via autofluorescence confocal microscopy. Constructs were fabricated to a 200 µm height using 10 µm layer thicknesses and 5s exposures at 16 mW/cm². The hydrogel successfully resolved (D) the alternating spokes of the bicycle, establishing a minimum X/Y print resolution of ∼75 µm (scale bar = 750 µm); (E) the distinct right angles and varying dimensions of the “HI” beams with minimal rounding (scale bar = 250 µm); and (F) the continuous walls of the honeycomb structure without tearing (scale bar = 750 µm). (G) CAD rendering of the Vanderbilt University Medical Center (VUMC) logo. (H) Maximal Z-projection of the bioprinted VUMC logo encapsulated with normal human dermal fibroblasts (hNDFs) after 21 days of culture (scale bar = 1 mm). Fibroblasts extensively proliferated and remodeled the hydrogel, establishing an interconnected mesh network with active extracellular matrix deposition (ECM). Green = F-actin; Red = Collagen IV; Blue = nuclei (DAPI).

Across all three test geometries, the 4D hydrogel reproduced the corresponding CAD designs with high pattern fidelity (**Figure 3 A-F**). In the bicycle construct, the alternating positive and negative spoke features remained clearly separated with minimal bridging, demonstrating that closely spaced microfeatures could be resolved within a complex curved geometry. Distinct spatial features were maintained approximately 170 µm from the central hub, establishing a minimum lateral printing resolution of approximately 75 µm. The “HI” construct further demonstrated dimensional control across 100, 150, and 200 µm beam widths, with printed features retaining defined edges and minimal rounding despite height-to-width aspect ratios of 2:1, 1.5:1, and 1:1. At the larger construct scale, the honeycomb lattice maintained a continuous wall architecture without tearing, discontinuity, or loss of pattern continuity, indicating that the hydrogel withstood the cupping, suction, and blowout forces associated with repeated vertical movement during layer-by-layer DMD fabrication. Together, these acellular printing studies demonstrate that the 4D formulation supports both microscale feature resolution and construct-level geometric stability.

To determine whether high pattern fidelity was compatible with post-print cellular remodeling, hNDFs at a concentration of 2 × 10⁶ cells mL⁻¹ were encapsulated within the optimized 4D GelMA-Cad/gelatin bioink and printed into a larger VUMC logo geometry (**Figure 3G, H**). This experiment extended the acellular fidelity studies into a cell-laden construct, where the material must preserve the intended printed architecture while also permitting the embedded cells to reorganize their local microenvironment. After 21 days of culture, hNDFs remained distributed throughout the patterned construct and adopted an extensively spread morphology, forming dense, interconnected cellular networks within the hydrogel. The F-actin organization demonstrated robust cytoskeletal remodeling across the printed structure, indicating that the matrix permitted cell attachment, spreading, and multicellular connectivity after fabrication (**Figure 3 H**). Collagen IV staining further revealed active extracellular matrix deposition by the embedded fibroblasts, supporting the conclusion that the 4D hydrogel was not simply permissive to cell survival, but also to matrix remodeling and de novo ECM assembly (**Figure 3I**). This distinction is important because hydrogels optimized for high-resolution printing can preserve shape by remaining mechanically restrictive ^[57]^, which may limit the cellular remodeling needed for tissue maturation ^[58–60]^. In contrast, the GelMA-Cad/gelatin formulation maintained the geometric advantages required for DMD fabrication while transitioning into a matrix environment that supported fibroblast-mediated remodeling over long-term culture. Thus, the same formulation that enabled microscale print fidelity also permitted the cellular behaviors needed to begin replacing the initially printed hydrogel architecture with cell-organized matrix structure.

These acellular and cell-laden printing studies establish that the 4D hydrogel meets the two functional requirements that motivated its design: high-fidelity DMD patterning during fabrication and matrix permissiveness during culture. The CAD-based geometries demonstrated that the formulation could reproduce complex projected masks containing small negative spaces, variable beam widths, high-aspect-ratio features, and larger continuous-wall architectures while retaining pattern fidelity after post-print washing and early swelling equilibration. Extending this validation into a cell-laden VUMC logo construct showed that geometric stability was maintained in a biologically active setting, where embedded fibroblasts spread throughout the patterned matrix, organized their cytoskeleton, and deposited collagen IV over 21 days of culture (**Figure 3G, H)**. This combination of print fidelity and long-term cellular remodeling is essential for vascularized tissue fabrication, where the printed structure must persist long enough to guide construct formation while remaining permissive to matrix reorganization by endothelial cells, pericytes, and supporting fibroblasts. The 4D GelMA-Cad/gelatin bioink therefore provides a validated material platform for moving beyond single-cell-type patterning toward multicellular, lumen-forming vascular tissue models, beginning with retinal vascular tricultures designed to recapitulate key features of early iBRB-like assembly.

### 3.3 Retinal Vascular Tricultures Form Lumenized Networks within the 4D Matrix

Following validation of DMD pattern fidelity and fibroblast-mediated matrix remodeling, the 4D GelMA-Cad/gelatin hydrogel was evaluated for multicellular vascular assembly in a retinal barrier context. The iBRB is built around specialized retinal microvascular endothelium and its interactions with perivascular support cells and basement membrane-rich extracellular matrix; therefore, modeling early iBRB-like vascular organization requires more than endothelial cell survival alone. To capture key cellular components of this vascular interface, bioprinted retinal tricultures were generated using human primary retinal endothelial cells (hPRECs), human primary retinal pericytes (hPRPs), and normal human dermal fibroblasts (hNDFs) within 6 mm diameter, 200 µm-thick hydrogel discs (**Figure 4A–D**). The bioink was prepared at a total cell density of 6 × 10⁶ cells mL⁻¹ using a 4:1:1 endothelial:pericyte:fibroblast ratio, establishing a defined multicellular system in which retinal endothelial cells formed the primary vascular compartment, pericytes provided mural support, and fibroblasts contributed matrix-remodeling and ECM support. This model was designed to evaluate whether the 4D matrix could support the foundational cellular events of iBRB-like vascular assembly after printing, including retinal endothelial network formation, perivascular support-cell interactions, matrix remodeling, and lumenization.

**Figure 4.**
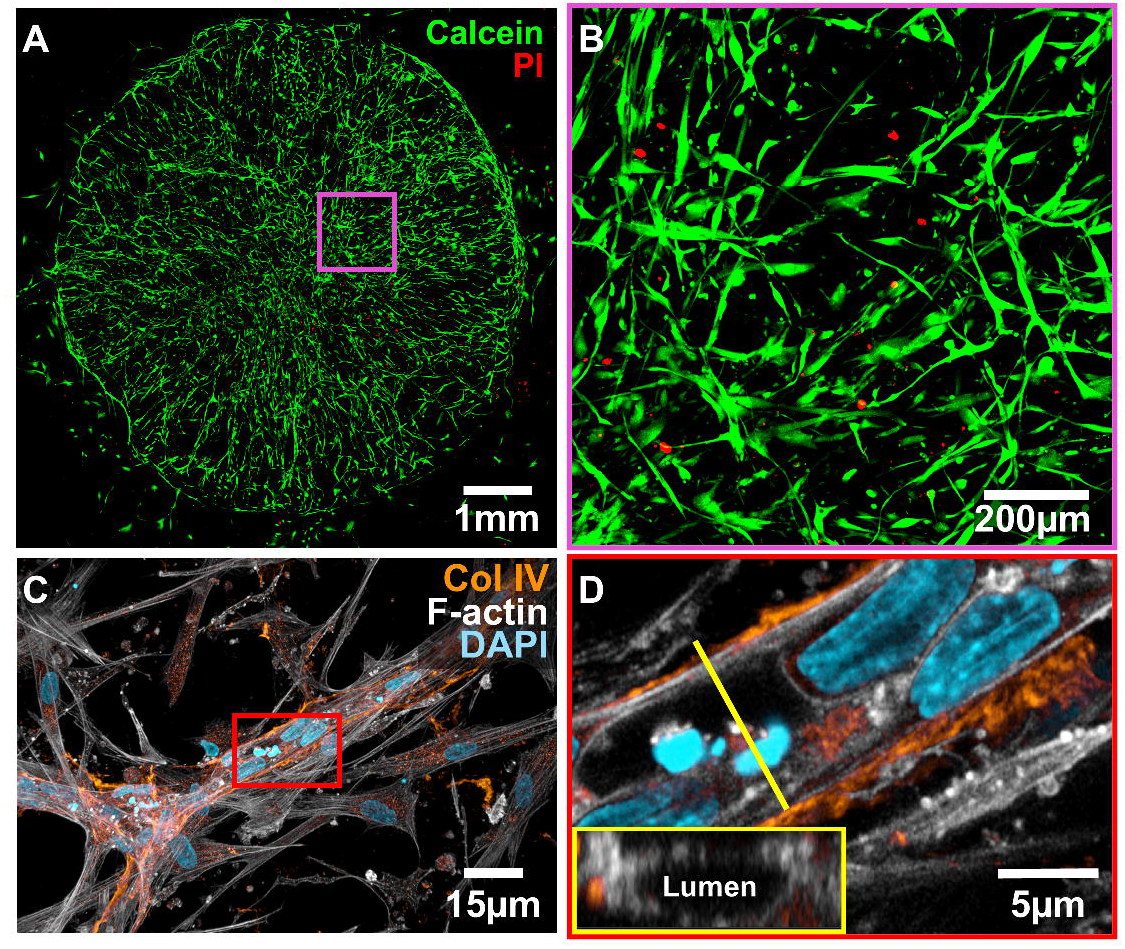
Self-assembly and vascular network formation of a 3D bioprinted *in vitro* inner blood-retinal barrier (iBRB) model. (A) Maximal Z-projection showing live/dead cellular viability within a bioprinted triculture construct after 7 days of culture. The 4D hydrogel (6mm diameter, 200µm thickness) encapsulated a mixture of human primary retinal endothelial cells (hPRECs), human primary retinal pericytes (hPRPs), and normal human dermal fibroblasts (hNDFs), demonstrating robust cytocompatibility with 99.78% live cells and 0.22% dead cells (scale bar = 1mm). (B) High-magnification view of the live/dead stain detailing extensive cellular spreading and the rapid formation of dense, interconnected mesh networks throughout the hydrogel (scale bar = 200µm). (C) Immunofluorescence microscopy of the triculture construct at Day 7, revealing active cellular remodeling of the matrix and early stages of vascularization (scale bar = 15µm). (D) High-magnification and orthogonal cross-sectional views highlighting distinct lumen formation within the self-assembled vascular structures, supported by localized collagen IV deposition from pericytes and fibroblasts (scale bar = 5µm). For (A) and (B): Green = Live cells (Calcein-AM); Red = Dead cells (Propidium Iodide). For (C) and (D): Orange = Collagen IV; White= F-actin; Cyan = nuclei (DAPI).

After 7 days of culture, the retinal tricultures demonstrated high cytocompatibility within the 4D matrix. Live/dead staining with calcein-AM and propidium iodide showed 99.78% live cells and 0.22% dead cells, indicating that the hydrogel formulation and DMD fabrication conditions were well tolerated by the encapsulated vascular cell populations (**Figure 4A; Figure S6A, B)**. High-magnification imaging further showed that the cells were not maintained as isolated rounded populations, but instead spread extensively through the construct and formed dense, interconnected multicellular mesh-like networks (**Figure 4B**). This early network formation is consistent with the mechanical behavior established in Section 3.1, where gelatin-mediated softening and pore opening generated a more permissive matrix state during culture. The combination of high viability, extensive cell spreading, and dense multicellular connectivity indicates that the 4D hydrogel provides a cytocompatible and physically permissive environment for retinal vascular triculture assembly.

Day 7 immunofluorescence demonstrated that the retinal tricultures advanced from early cell spreading into coordinated matrix remodeling and lumenized vascular organization. Collagen IV is localized around multicellular network structures, indicating deposition of vascular-associated extracellular matrix within the printed hydrogel (**Figure 4C, D)**. This collagen IV-rich organization reflects remodeling by the triculture as an integrated system, with fibroblasts and pericytes contributing supportive matrix-remodeling functions within the developing endothelial network. These findings are particularly relevant to the GelMA-Cad design, which distinguishes the matrix from a purely structural GelMA/gelatin scaffold by introducing covalently incorporated N-cadherin-derived adhesion cues relevant to endothelial–pericyte communication and multicellular vascular organization ^[50]^. In this context, the N-cadherin-mimetic component provides a bioinstructive adhesion cue aligned with the coordinated cell–cell interactions required for vascular assembly. Within this remodeled hydrogel, confocal imaging revealed organized F-actin-positive multicellular structures, and orthogonal views confirmed distinct lumen-containing vascular networks (**Figure 4C, D)**. The observed lumens were approximately 5 µm in diameter, placing them within a capillary-scale range relevant to human retinal microvascular architecture.

These results are consistent with prior work demonstrating that human vascular tissues can undergo lumenization and organize into concentric vascular architectures within permissive hydrogel environments ^[50]^. The collagen IV-rich matrix organization, F-actin-positive cellular architecture, and capillary-scale lumen formation support the conclusion that the 4D matrix provides both the physical and bioinstructive environment needed for early iBRB-like vascular organization.

The retinal triculture experiments provide the first biological validation that the 4D matrix can translate print-stage material properties into coordinated vascular tissue assembly. Within seven days, retinal endothelial cells, pericytes, and fibroblasts remained highly viable, spread extensively through the printed construct, and reorganized into dense multicellular networks enriched with collagen IV. The emergence of capillary-scale lumens within these remodeled structures is particularly important because it links the material’s engineered permissiveness to a defining feature of vascular organization. Gelatin-mediated softening and pore opening created a matrix environment that supported cellular reorganization, while the covalently incorporated N-cadherin-derived peptide provided bioinstructive adhesion cues relevant to endothelial–pericyte interactions. In this retinal barrier context, the same material features that enabled DMD print fidelity also supported multicellular remodeling, vascular-associated matrix deposition, and lumen-forming network assembly, establishing the 4D hydrogel as a platform for early iBRB-like vascular organization.

### 3.4 Bioprinted Pulmonary Vascular-Alveolar Constructs Develop Lumenized Vascular Beds and AT1-like Epithelial States in the 4D Hydrogel

The retinal vascular studies established that the 4D GelMA-Cad/gelatin hydrogel could support viable multicellular remodeling and lumenized microvascular assembly, motivating a more complex test of whether the same material strategy could organize a vascular–alveolar tissue interface (**Figure 4 A-D; Figure S6A, B)**. Pulmonary vascular–alveolar assembly places a higher biological demand on the matrix because endothelial lumen formation, fibroblast-mediated remodeling, epithelial organoid incorporation, and local vascular–epithelial organization must occur within a shared three-dimensional environment.

To build this system, pulmonary vascular beds were generated by bioprinting human lung microvascular endothelial cells (HULECs) and human lung fibroblasts (HLFs) within 6 mm diameter, 150 µm-thick hydrogel discs with a cell density of 5 × 10⁶ cells mL⁻¹ at a 7:1 endothelial:fibroblast ratio. By day 7, the HULEC/HLF vascular bed constructs showed extensive multicellular organization throughout the printed matrix, with endothelial and fibroblast populations assembling into interconnected vascular structures within the engineered hydrogel (**Figure 5A, B)**. VE-cadherin staining revealed endothelial junctional organization along these networks, while F-actin highlighted cytoskeletal spreading and multicellular connectivity across the developing vascular bed. PODXL expression was especially informative in this setting because luminal/apical organization is a defining feature of vascular lumen formation and endothelial polarization within three-dimensional tissues ^[61–63]^. In the pulmonary constructs, PODXL localized to endothelial structures that contained visible lumen-like spaces by orthogonal confocal imaging, linking this apical-associated marker to a clear structural feature of the vascular bed rather than to endothelial identity alone. This combination of VE-cadherin-positive endothelial organization, PODXL-associated luminal patterning, and cross-sectional evidence of lumen formation supports the conclusion that the printed matrix permits pulmonary endothelial cells and fibroblasts to assemble into lumenized vascular beds within the 4D hydrogel environment.

**Figure 5.**
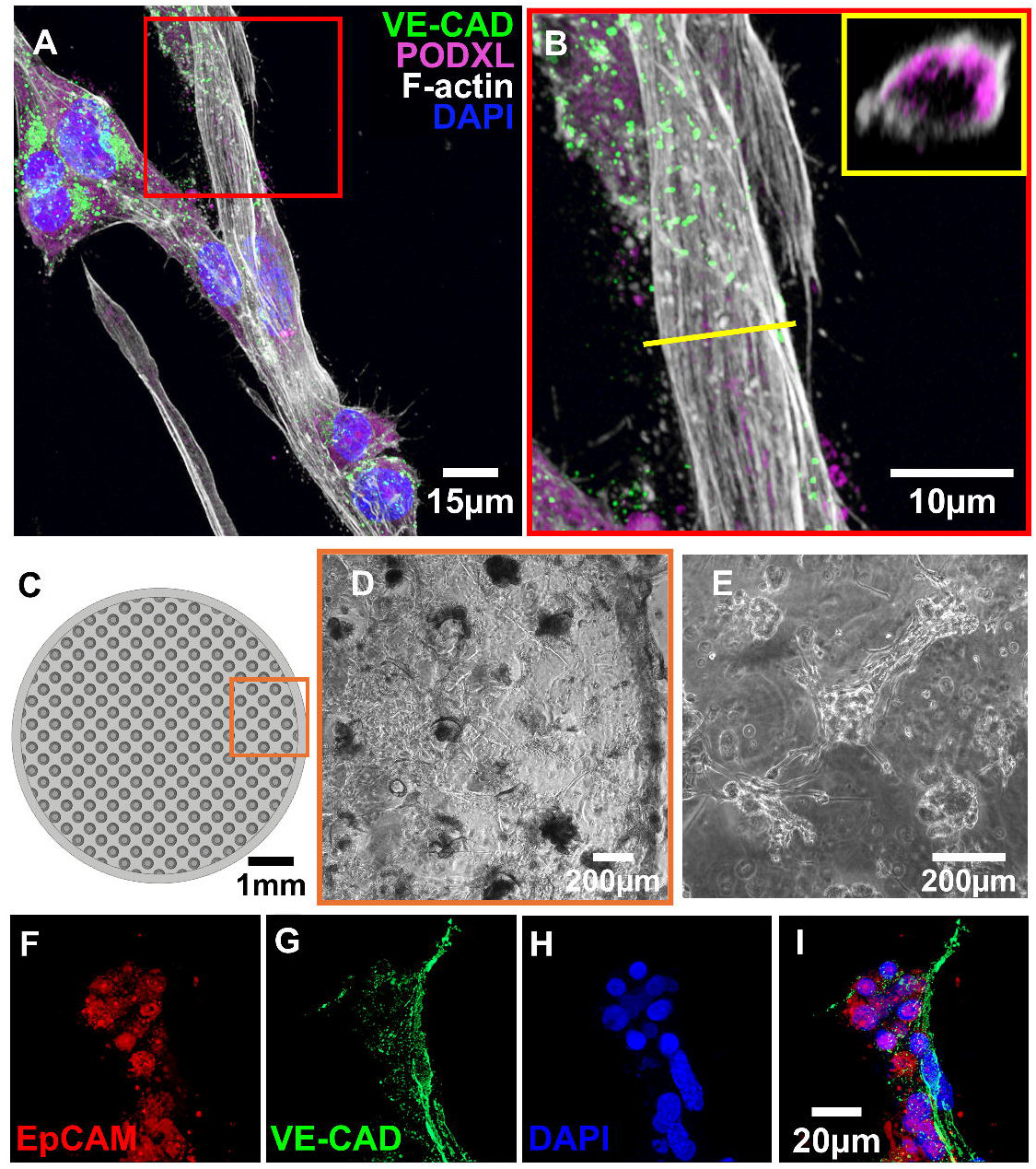
4D hydrogel-embedded lung vascular constructs support lumen formation and alveolar organoid integration . (A) Immunofluorescence staining of a lung vascular bed construct at Day 7 after bioprinting. Constructs contained human lung microvascular endothelial cells (HULECs) and human primary lung fibroblasts (HLFs) at a cell density of 5 × 10⁶ cells mL⁻¹ and a 7:1 ratio within 4D GelMA-Cad/gelatin hydrogels measuring 6 mm in diameter and 150 µm in thickness. VE-cadherin is shown in green, PODXL in magenta, F-actin in white, and DAPI in blue. Scale bar = 15 µm. (B) High-magnification view of the boxed region in (A), with orthogonal cross-sectional visualization showing lumen formation and PODXL-associated luminal/apical organization (yellow inset). VE-cadherin is shown in green, PODXL in magenta, F-actin in white, and DAPI in blue. Scale bar = 10 µm. (C) Top view of microwell CAD design for trapping lung organoids with embedded lung vascular bed. (D) Brightfield image of the bioprinted construct 24 hours after placing lung organoids on the construct (5x magnification). (E) Brightfield image 24 hours after adding organoids with the vascular bed integrating with the organoids (10x magnification). (F–I) Immunofluorescence staining of day 14 lung vascular bed constructs printed with a cell density of 4.5 × 10⁶ cells mL⁻¹ and an 8:1 ratio of HULEC/HLF with organoids seeded 4 days post bioprinting. (F) EpCAM, (G) VE-cadherin, (H) DAPI, and (I) merged image showing EpCAM-positive epithelial structures associated with VE-cadherin-positive vascular networks. Scale bar = 20 µm. Data from 3 independent experiments.

To incorporate the alveolar organoids into the vascular bed, a microwell design was implemented to physically position the epithelial compartment within the developing pulmonary vascular matrix. The 6mm diameter disc features a 100µm-tall base to serve as the foundation for the pulmonary vascular bed beneath an array of 200µm-tall microwells, resulting in a 300 µm-tall hydrogel construct with evenly spaced 250 µm-wide microwells that cradle the organoids allowing them to incorporate into both the hydrogel and pulmonary vascular bed upon seeding (**Figure 5C; Figure S7A)**. To bioprint the pulmonary vascular lung construct, HULEC and HLF were prepared in a 16:1 ratio and cultured for 5 days to establish the pulmonary vascular bed prior to seeding alveolar organoids dropwise into the microwells. This staged design created a vascularized hydrogel environment for evaluating lumenized pulmonary vascular bed formation, epithelial organoid incorporation, and local vascular–alveolar organization within the same engineered construct, while allowing the endothelial–fibroblast network to establish before introduction of the alveolar epithelial compartment. The pulmonary vascular constructs were then imaged via brightfield microscopy one day post organoid seeding and demonstrated preserved microwell architecture, retention of organoid positioning, clear incorporation of the organoids into the pulmonary vascular bed within 24 h of culture (**Figure 5D, E)**. To further interrogate the spatial tissue organization of this pulmonary construct after long-term culture, HULECs and HLF were bioprinted with a cell density of 4.5 × 10⁶ cells mL⁻¹ in a ratio of 8:1 and allowed to grow for 4 days prior to seeding alveolar organoids and maintained for an additional 10 days of culture. EpCAM-positive epithelial structures were retained within the construct and were positioned in close proximity to VE-cadherin-positive vascular networks (**Figure 5F–I**). This local association between epithelial organoids and endothelial structures is an important outcome of the model, as vascular–alveolar tissue organization depends on spatial coordination between the epithelial and vascular compartments. Rather than serving only as a geometric holding feature, the microwell architecture enabled epithelial organoids to remain localized within the vascularized construct while endothelial networks developed nearby within the surrounding matrix **(Supplementary Video 1)**. Together, these imaging results establish the pulmonary construct as a multicellular vascular–epithelial system in which lumenized endothelial networks, PODXL-associated luminal/apical organization, and EpCAM-positive epithelial organoid incorporation occur within the same vascularized hydrogel environment.

With the multi-compartment pulmonary construct established, scRNA-seq was used to determine how the vascularized hydrogel environment shaped the cellular state composition and the spatial juxtaposition of the different cell types were altered through this designed spatial juxtaposition. Cells were isolated from the co-cultured print at day 17 post bioprinting. This analysis included 10,696 cells collected from input/control cell populations and from cells isolated after culture within the bioprinted pulmonary constructs, allowing direct comparison between the starting populations and the co-cultured printed tissue environment. Expected endothelial, epithelial, and fibroblast compartments were identified and clearly separated within the integrated dataset (**Figure 6A, B and Figure S7B)**. Compartment annotation was derived from canonical markers, including epithelial, endothelial, and fibroblast-associated genes that defined each major population within the printed construct and input cell populations (**Figure 6C**). This scRNA-seq framework therefore provided the basis for evaluating whether the structural organization observed in Figure 5 was accompanied by compartment-specific transcriptional changes within the pulmonary vascular– alveolar model.

**Figure 6.**
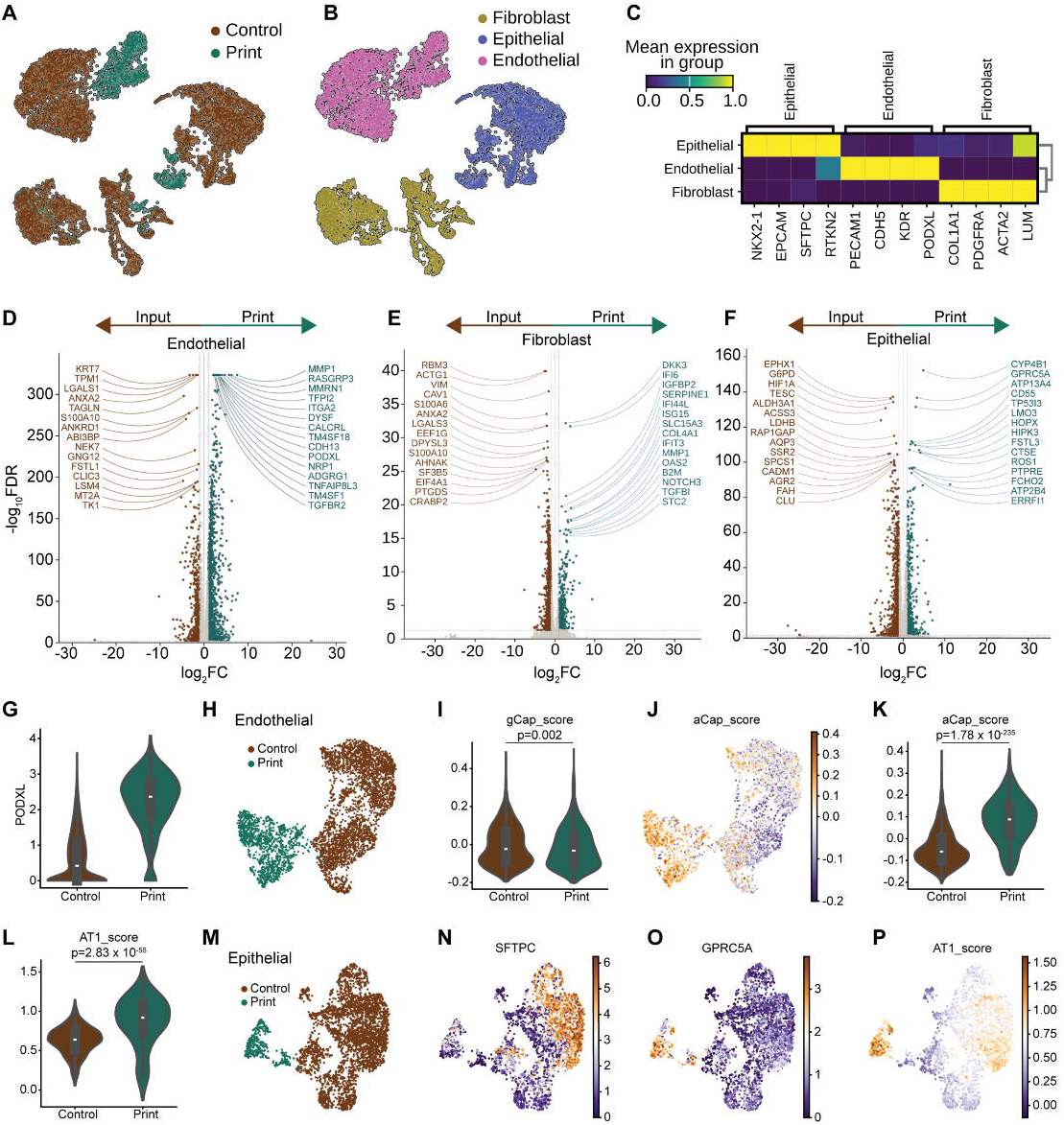
Bioprinted pulmonary constructs develop endothelial and epithelial transcriptional programs associated with vascular–alveolar remodeling. UMAP visualization of scRNA-seq data from 10,696 cells derived from input cell populations and cells isolated from bioprinted pulmonary constructs following 17 days of culture. Cells are colored by experimental conditions in (A) and by annotated cell compartment in (B). (C) Matrix plot of canonical marker genes used to annotate epithelial, endothelial, and fibroblast compartments. (D-F) Volcano plots showing differentially expressed genes within isolated (D) endothelial, (E) fibroblast, and (F) epithelial compartments. Differential expression testing was performed within each annotated compartment using Wilcoxon rank-sum analysis. (G) Violin plot showing expression of PODXL within the endothelial subset. (H) UMAP projection comprised of scRNA-seq data from endothelial cells pooled from bioprinted pulmonary construct and control. (I) gCap endothelial signature scores derived from gene sets reported by Gillich et al. **^[67]^**, Travaglini et al. **^[69]^**, and Schupp et al. **^[70]^**. (J) UMAP overlay of aCap score on endothelial cell compartment. (K) aCap endothelial signature scores. (L) AT1 epithelial signature score derived from canonical atlas-based AT1 epithelial markers reported by Burgess et al. **^[68]^**, showing increased AT1-associated transcriptional enrichment within the epithelial compartment following culture in bioprinted pulmonary constructs. (M) UMAP projection comprised of scRNA-seq data from epithelial cells pooled from bioprinted pulmonary construct and control. UMAP overlays showing epithelial transcriptional patterning across the organoid-derived compartment, including (N) SFTPC, (O) GPRC5A, and (P) AT1_score.

Compartment-resolved differential expression analysis showed that the printed pulmonary environment was associated with distinct differential expression within endothelial, fibroblast, and epithelial subsets (**Figure 6D–F**). Deeper analysis of the endothelial subset supported both the polarization observed in whole-mount print imaging as well as enhancement of similarity to the specific capillary network at the blood-gas barrier where endothelium and epithelium interface. PODXL expression increased in the printed condition (**Figure 6G**), directly aligning with the PODXL-associated lumen structures observed by immunofluorescence in **Figure 5**. This agreement between protein-level imaging and single-cell transcriptomic analysis strengthens the interpretation that endothelial cells within the 4D matrix acquire luminal/apical vascular-associated features as they organize into three-dimensional vascular beds. Within the lung, endothelial gCap (general capillary) serve as vasomotor and stem-cell niche functions, while aCap (alveolar capillary) are lung-specific and the terminally differentiated endothelial surface of the blood-gas barrier. To date, there is minimal reported *ex vivo* exploration of the juxtaposition of primary endothelium-epithelium in this context, though there has been substantive iPSC related work ^[64–66]^. Endothelial gCap (general capillary) and aCap (alveolar capillary) scores were calculated from published lung capillary endothelial gene sets ^[67]^, revealing significant enhancement of aCap scores in the print, with essentially unchanged gCap scores (**Figure 6H-K**). This result extends the structural vascular data in **Figure 5** by showing that the lumenized endothelial networks were accompanied by single-cell transcriptional features associated with lung capillary endothelium and significantly enhanced signatures associated with terminally differentiated alveolar endothelium.

The epithelial compartment showed a complementary blood-gas barrier-like differentiation response with a transition from facultative progenitor Alveolar Type 2 (AT2)-like cells toward a more terminally differentiated AT1-like cell state within the print. CYP4B1, a marker for AT1 cells, expression increased within the epithelial subset after culture in the printed construct **(Figure S7C)** as did numerous other AT1-enriched genes with substantial specificity (e.g. ATP13A4, CD55, GPRC5A, AGER, HOPX, CLIIC3, CAV1) (**Figure 6F and Figure S7D-G)**. This strongly differs from input primary alveolar organoids cultured in domed Matrigel (**Figure 6F).** This finding is important because the epithelial organoids entered the system with an AT2-like/SPC-positive identity, and CYP4B1/GPRC5A/CD55 differential upregulation in the print strongly supports an AT1-like reprogramming. The AT1 signature score (derived from atlas-level and functional integration ^[68]^) was significantly increased (p= 2.83x10 ^-56^) in the printed condition (**Figure 6L**), supporting enrichment of an AT1-like epithelial feature plots further clarified how this transition was distributed across the epithelial manifold: SFTPC expression highlights original AT2/SPC-positive alveolar epithelial identity, while localized enrichment of AT1_score and GPRC5A identified regions acquiring AT1-like cell state (**Figure 6M-P**). This combination is particularly important because the epithelial organoids were introduced into the construct as an AT2-like population, yet culture within the vascularized bioprinted environment was associated with the emergence of AT1-like epithelial states. Together, the endothelial capillary signatures and epithelial AT1-like features suggest that the pulmonary construct supports coordinated and highly physiologically convergent transcriptional signatures found at the blood-gas barrier within the same engineered tissue environment.

Together, these findings demonstrate that the 4D bioink platform can support multicellular pulmonary vascular–alveolar tissue assembly in which vascular lumen formation and epithelial cell-state transition occur within a shared, remodeling hydrogel environment.

## 4. Conclusion

In this study, we establish a temporally tunable, bioinstructive 4D GelMA-Cad/gelatin bioink for high-resolution DMD bioprinting of vascularized soft tissue models. This platform addresses a fundamental materials limitation in light-based bioprinting: the need to preserve microscale architecture during fabrication while enabling the softer, remodeling-permissive matrix environment required for multicellular vascular organization after printing. By combining a photocrosslinked GelMA-Cad scaffold with covalently incorporated N-cadherin-derived adhesion cues and a physically entrapped gelatin phase, the bioink separates print-stage stability from post-print matrix mechanics. A central advantage of this system is its simplicity and scalability: temporal softening is built directly into the GelMA-Cad/gelatin formulation rather than through synthetic polymer networks, multistep crosslinking schemes, or secondary degradation process. The resulting hydrogel maintains patterned fidelity during DMD fabrication, then undergoes time-dependent softening and microstructural opening as gelatin diffuses from the scaffold without requiring bulk degradation of the primary GelMA-Cad network. This transition supports both the architectural demands of high-resolution printing and the biological demands of cell spreading, cytoskeletal organization, extracellular matrix deposition, and lumen-forming vascular remodeling. In this way, the material does more than preserve the printed shape; it creates a dynamic matrix environment that allows initially fabricated structures to become biologically organized tissues, as demonstrated across two biologically distinct tissue contexts: retinal iBRB-like vascular assembly and pulmonary vascular–alveolar modeling.

Beyond material validation, this strategy provides a versatile framework for engineering multicellular vascular interfaces across distinct soft tissue contexts. In retinal vascular tricultures, the 4D matrix supported robust cell viability, collagen IV-rich network remodeling, and capillary-scale lumen formation, establishing a path toward bioprinted models of early iBRB-like assembly.

In pulmonary vascular–alveolar constructs, the platform supported lumenized endothelial beds, PODXL-associated luminal/apical organization, microwell-guided alveolar organoid incorporation, and spatial association between epithelial structures and vascular networks. Single-cell RNA-sequencing further demonstrated that the printed pulmonary environment maintained endothelial, fibroblast, and epithelial compartments while promoting lung capillary-associated endothelial features and AT1-like epithelial states in tight spatial juxtaposition with highly physiologic transcriptional features not yet achieved with primary cell inputs from humans. Looking forward, the modularity of this approach should enable tissue-specific tuning of matrix softening, incorporation of additional vascular, stromal, epithelial, immune, or disease-relevant cell populations, and integration with perfusion, injury, disease modeling, and therapeutic screening workflows. By uniting high-resolution DMD patterning with time-dependent matrix remodeling, this work establishes 4D hydrogel design as a practical framework for vascularized human tissue models and positions temporal matrix mechanics as a broadly applicable design principle for engineering complex multicellular tissues.

## Acknowledgements

Cryo-electron microscopy was performed in part through the use of the Vanderbilt Cell Imaging Shared Resource (supported by NIH grants CA68485, DK20593, DK58404, DK59637, EY08126, S10OD028704, and 1S10MH137068). Acquisition of the 400 MHz NMR spectrometer was supported by grant 1S10OD036222-01 from the National Institutes of Health. Nicole T. Marguerite acknowledges support from the National Science Foundation Graduate Research Fellowship Program. Any opinions, findings, and conclusions or recommendations expressed in this material are those of the author(s) and do not necessarily reflect the views of the National Science Foundation. Jonathan A. Kropski acknowledges support from P01HL172729 (JAK). A. Scott McCall acknowledges support from the Pulmonary Fibrosis Foundation, Francis Family Foundation, Vanderbilt Faculty Research Scholars (VFRS), K08HL175235, University of Colorado School of Medicine Lung Fibrosis Center. Brian O’Grady acknowledges support from the National Institute of Neurological Disorders and Stroke (NINDS) of the National Institutes of Health under Award Numbers K99NS133399 and R00NS133399.

## Data Availability Statement

The data and supporting CAD designs that support the findings of this study are available from the corresponding author upon reasonable request.

---

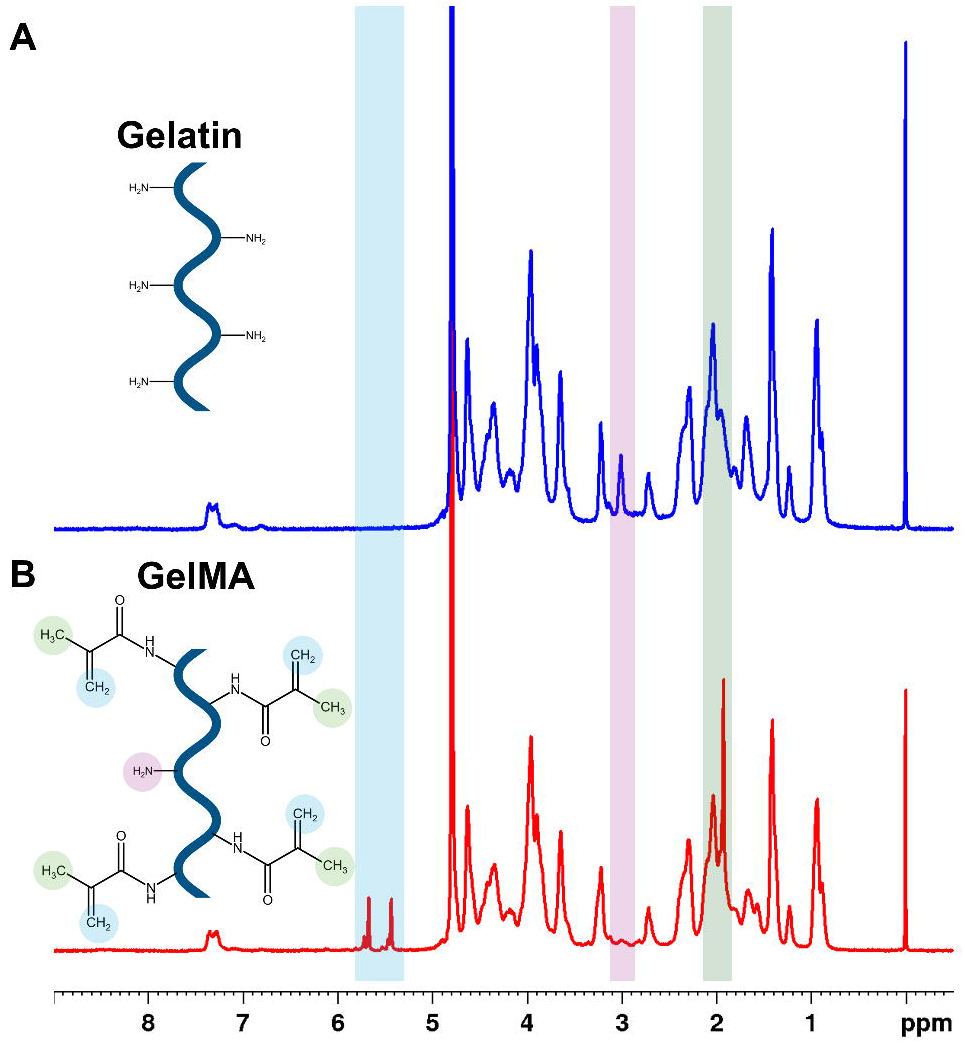

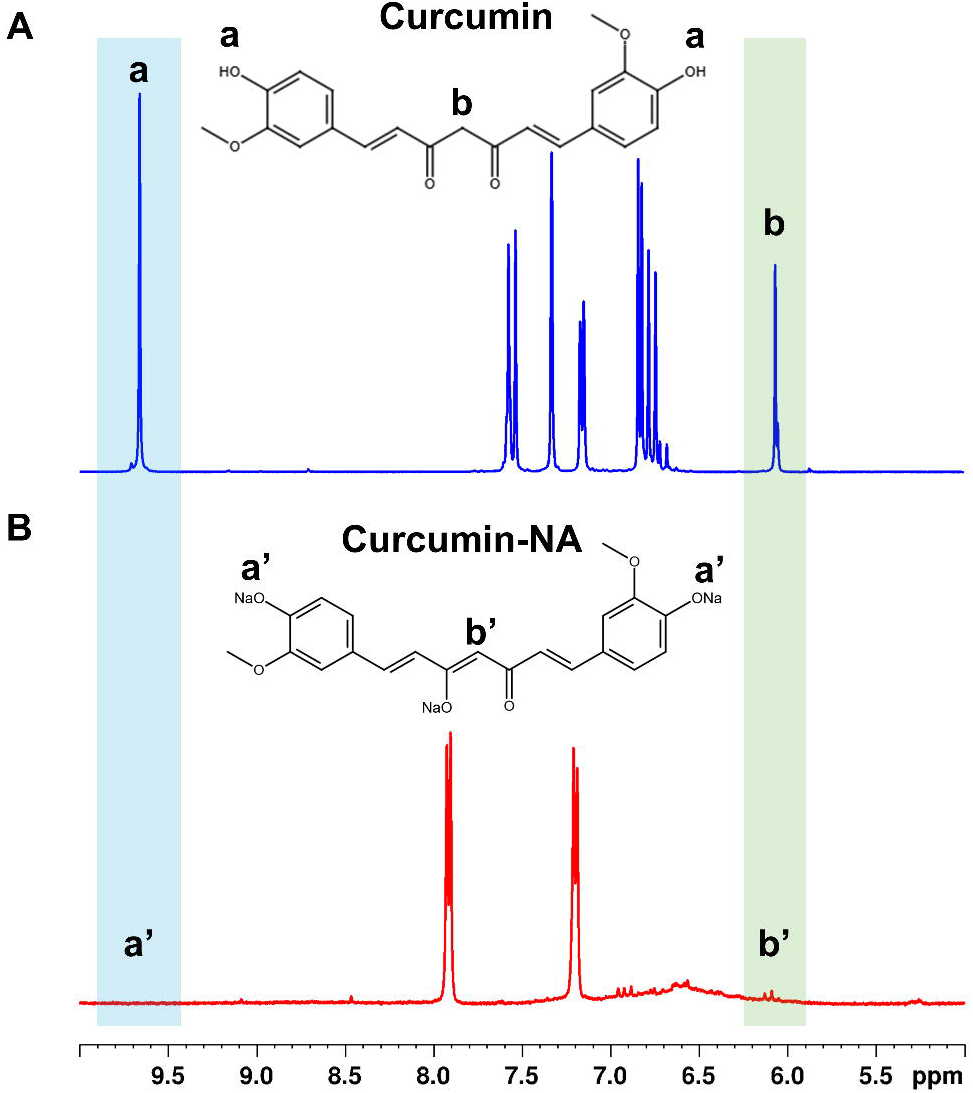

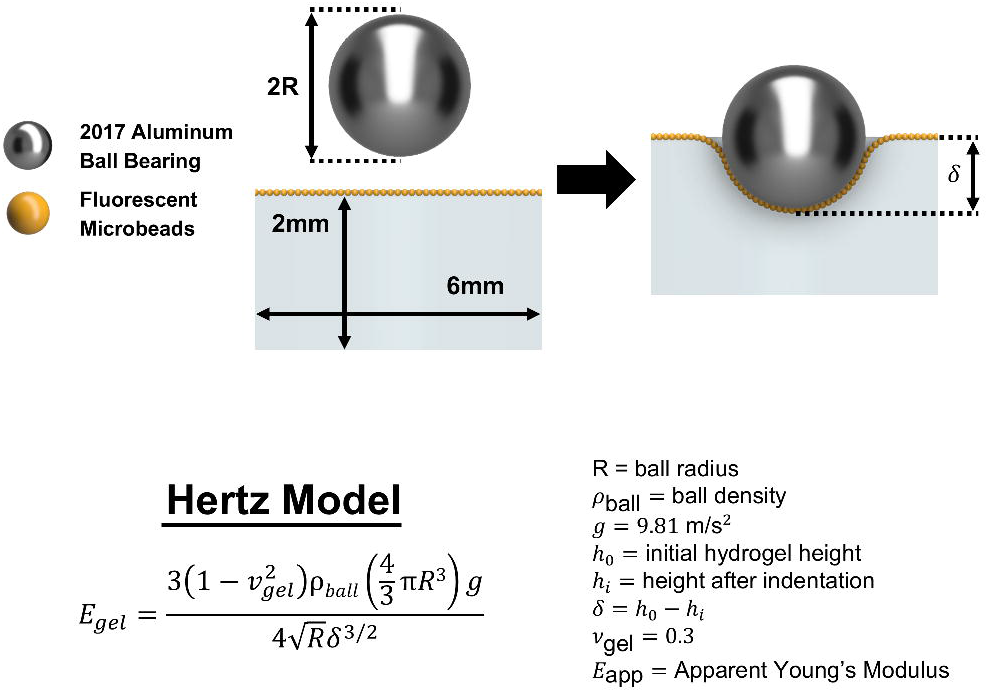

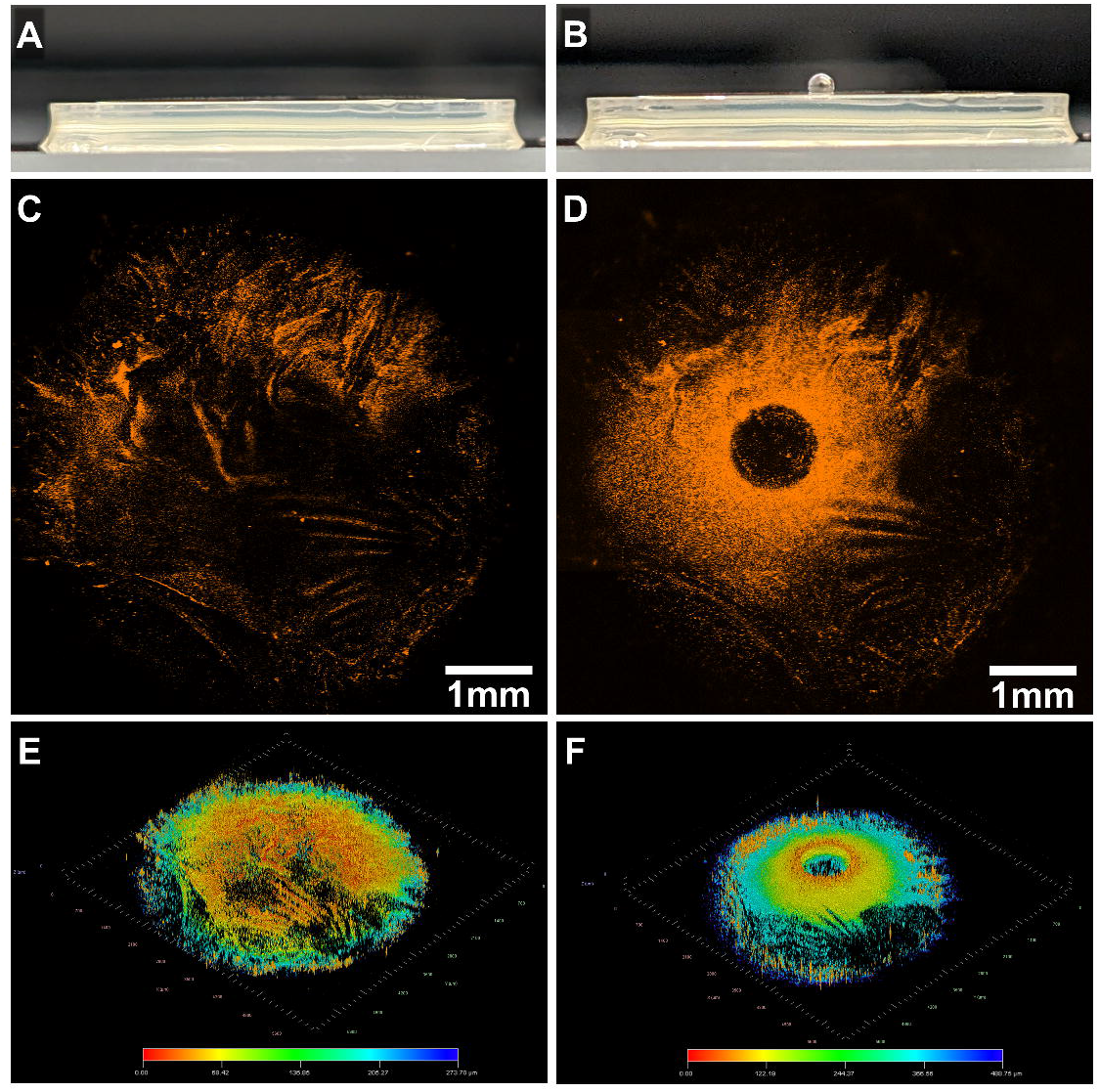

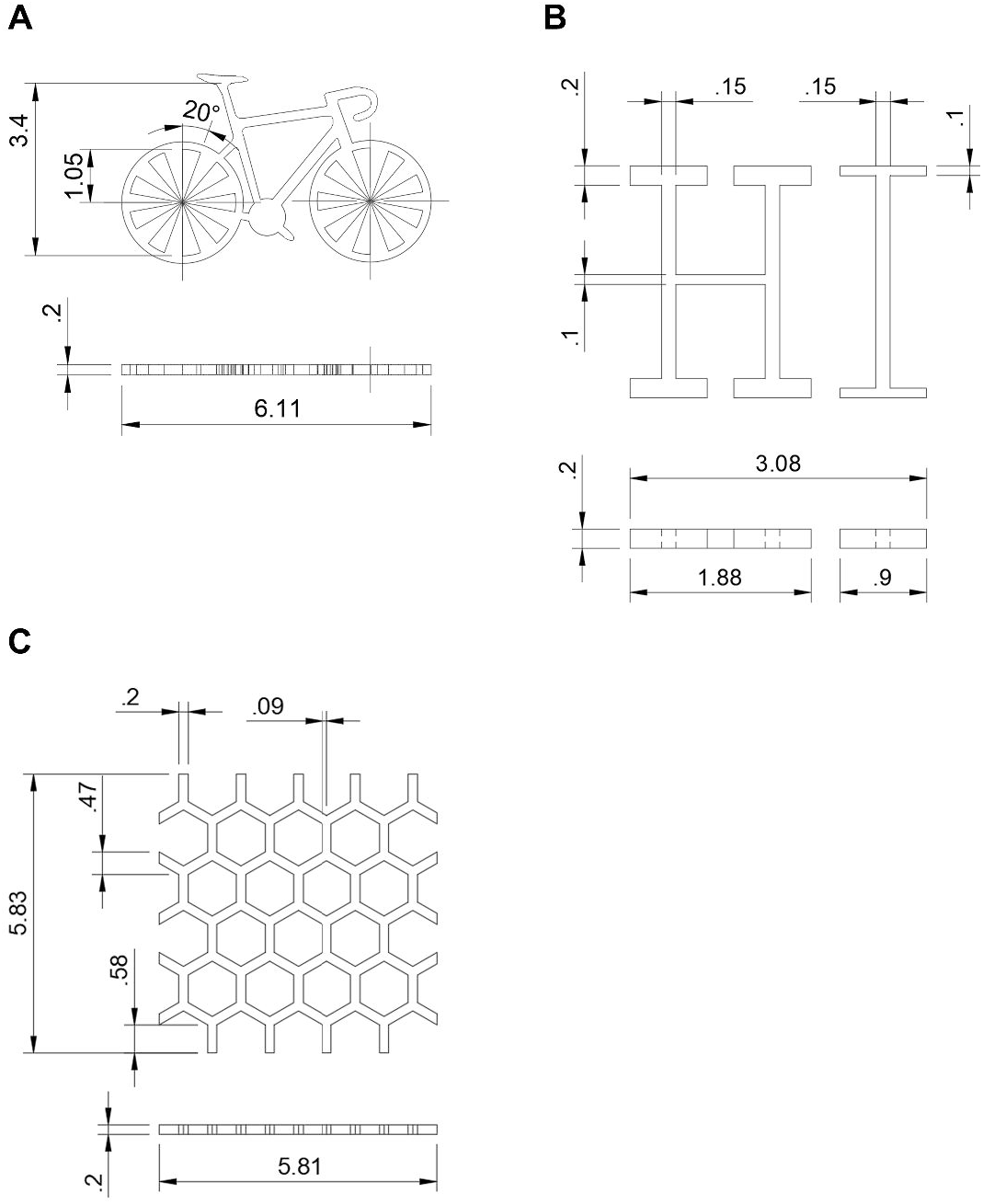

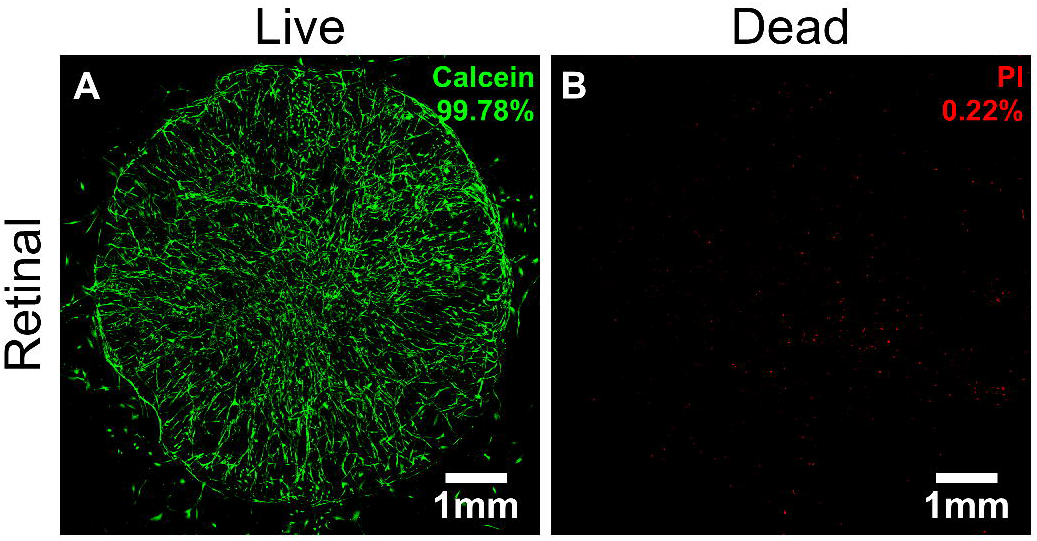

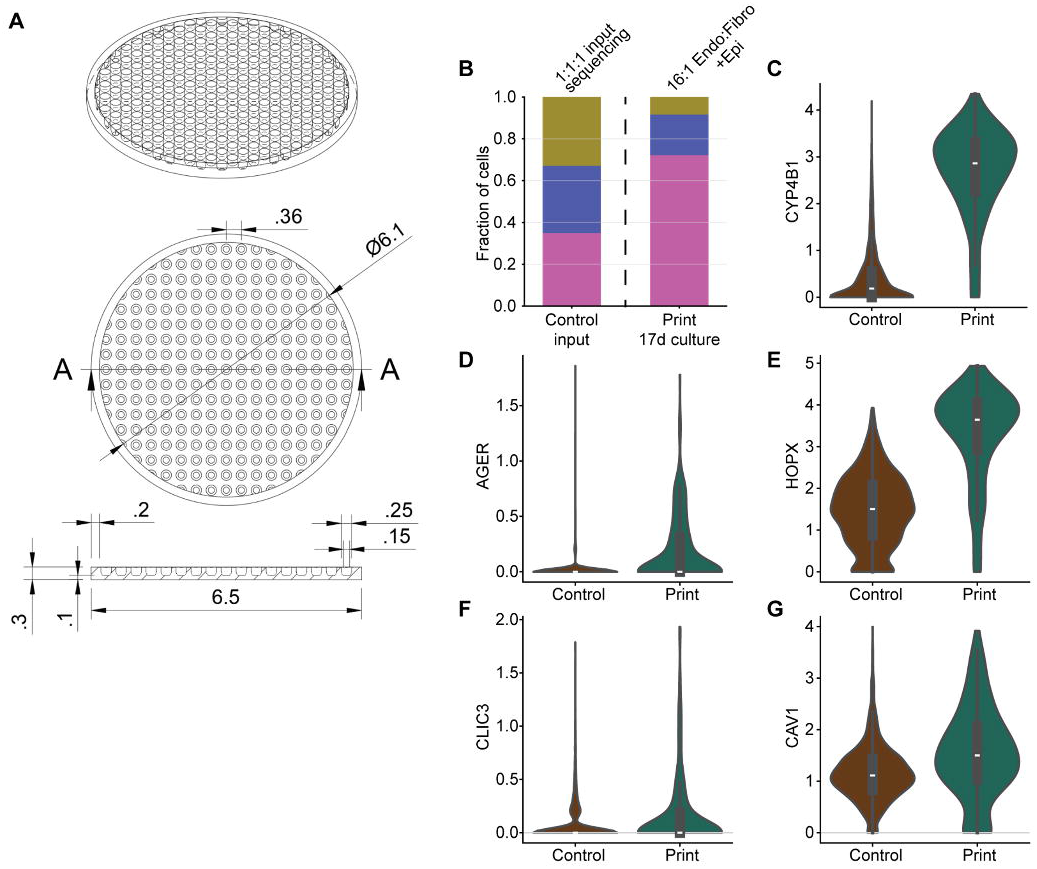

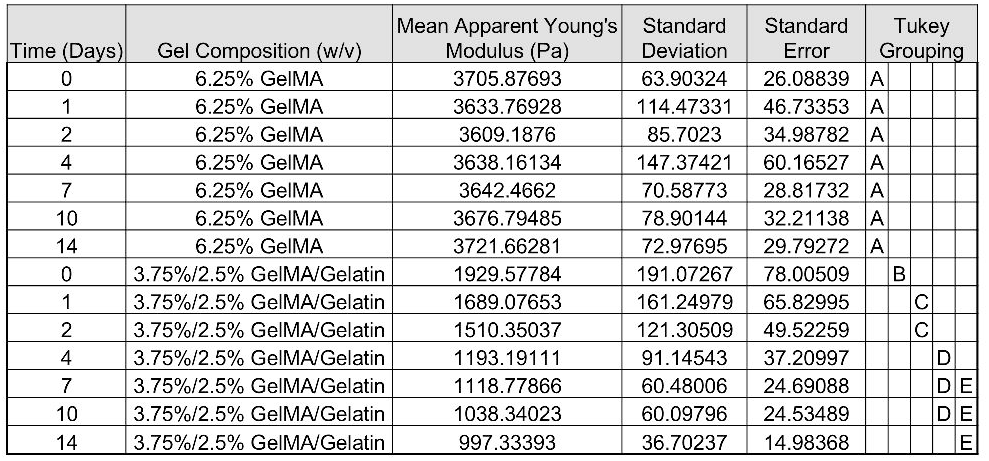

