## Supporting Information for "A 4D GelMA-Cad/Gelatin Hydrogel for High-Resolution DMD Bioprinting of Vascularized Soft Tissue Models"

**This Word document includes**:

Supplementary Figures: S1 to S7

Table S1

**Other supplementary materials for this manuscript include the following:**

Movie S1

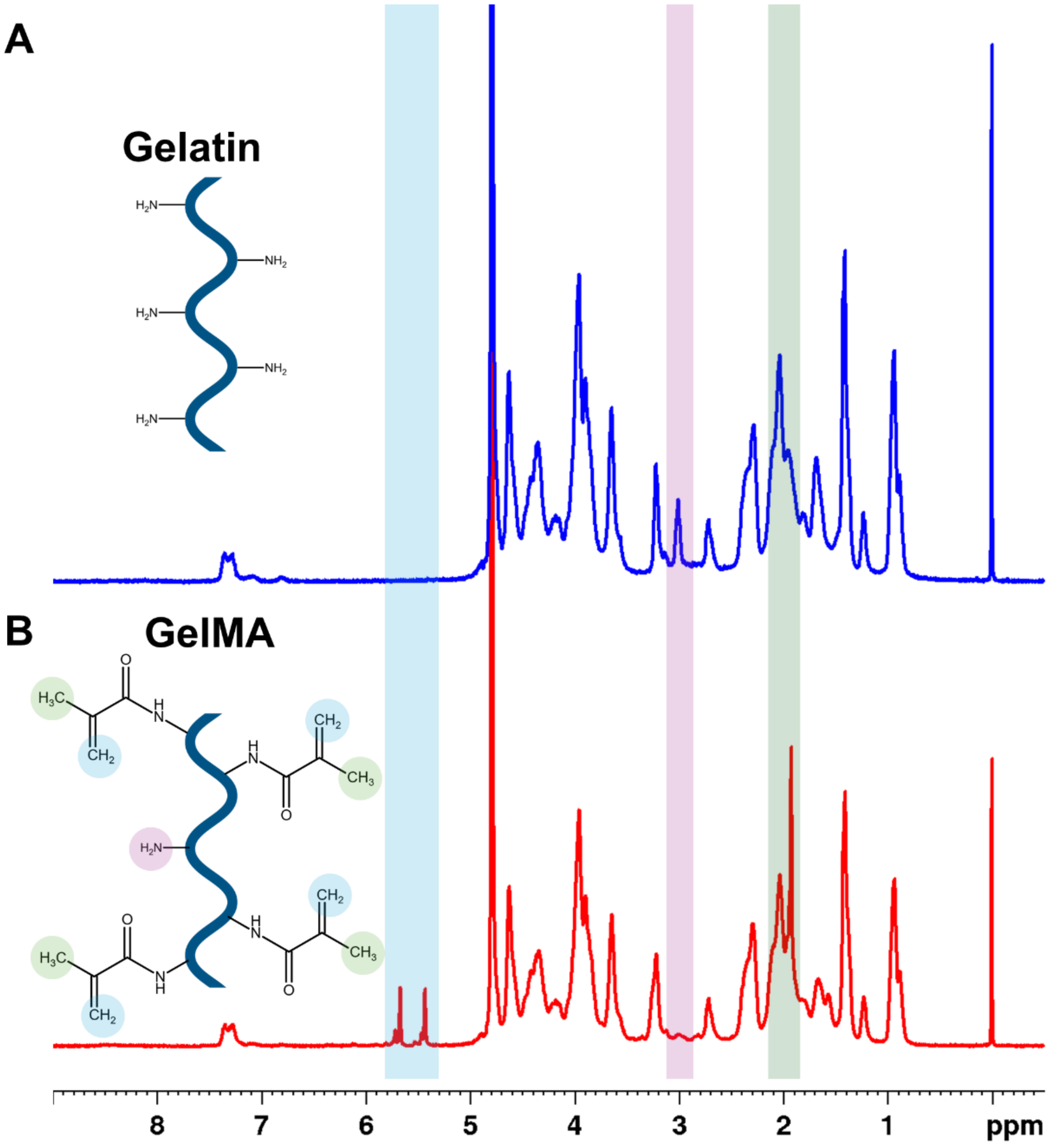


**Figure S1.** Chemical characterization of gelatin methacryloyl (GelMA) synthesis $^{1}$H-NMR spectroscopy. $^{1}$H-NMR spectra of (A) unmodified Type B gelatin and (B) synthesized GelMA, with corresponding chemical structures illustrating the methacryloyl substitution. Successful functionalization is confirmed by the appearance of new peaks between ~5.4–5.8 ppm corresponding to the vinylic protons of the grafted methacrylamide groups, and between ~1.8–

2.2 ppm corresponding to the methyl protons of the methacrylamide groups. The corresponding reduction of the peak between ~2.9–3.1 ppm, which represents the methylene protons of unreacted lysine and hydroxylysine residues, corresponds to a calculated degree of substitution of 92%**.** (Note: Colored highlight ranges for these specific peaks were applied to the spectra for visual clarity.)

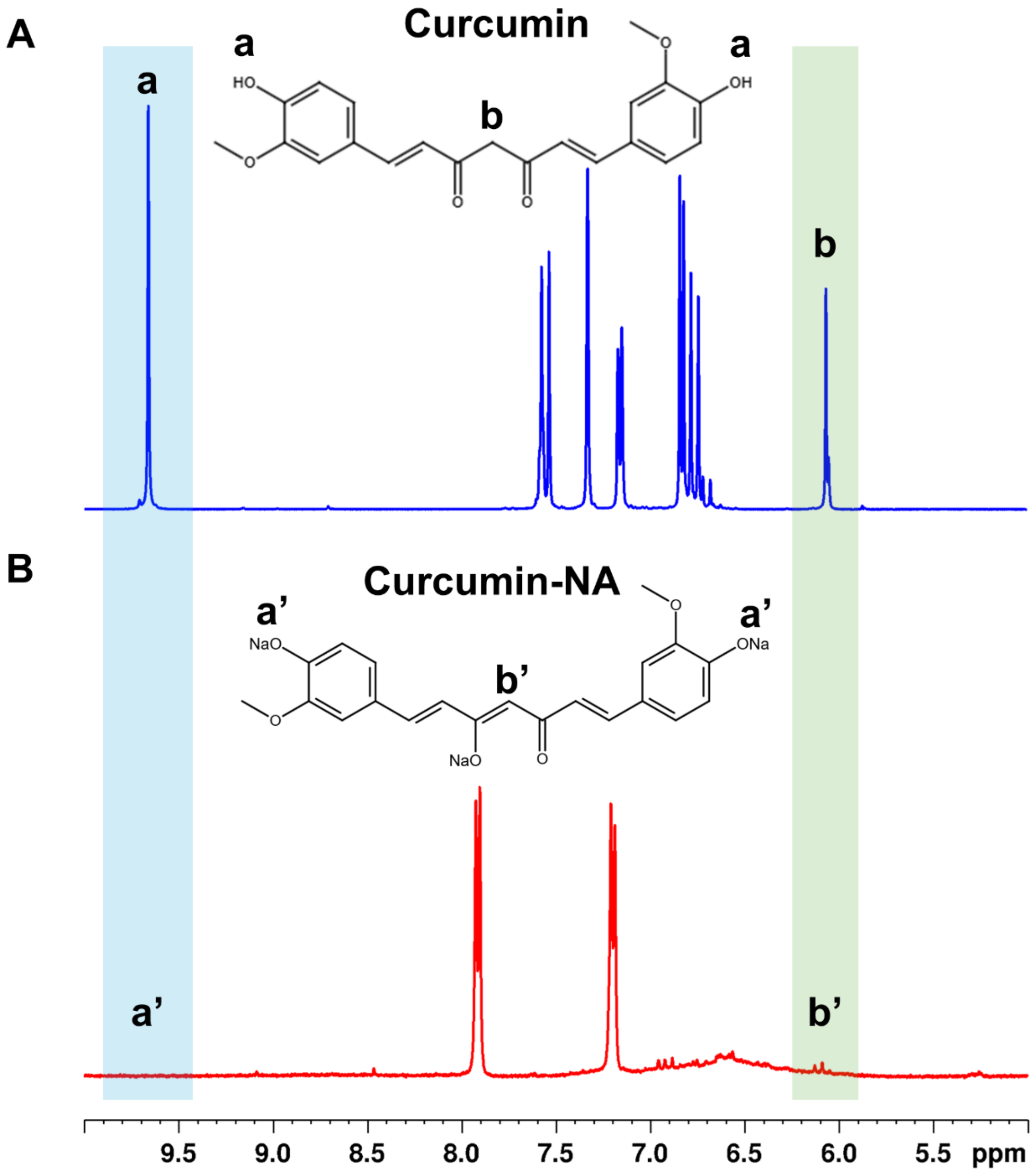


**Figure S2.** Chemical characterization of water-soluble Curcumin-NA synthesis via [1]H-NMRspectroscopy. [1]H-NMRspectra of (A) unmodified curcumin and (B) the synthesized Curcumin-NA salt. To overcome the inherent hydrophobicity of plain curcumin, sodium bicarbonate was used to substitute the protons of the phenolic hydroxyl groups with sodium ions yielding an aqueous-soluble photoabsorber as previously. Successful structural conversion is confirmed by the loss of resonance of the phenolic hydroxyl group at ~9.65 ppm (indicated by the structural transition from a to a'). Additionally, the modification influences the keto-enol tautomerism of the carbonyl backbone, evidenced by the chemical shift at ~6.1 ppm (indicated by the transition from b to b'). (Note: Colored highlights representing these specific shifts were applied to the spectra for visual clarity.)

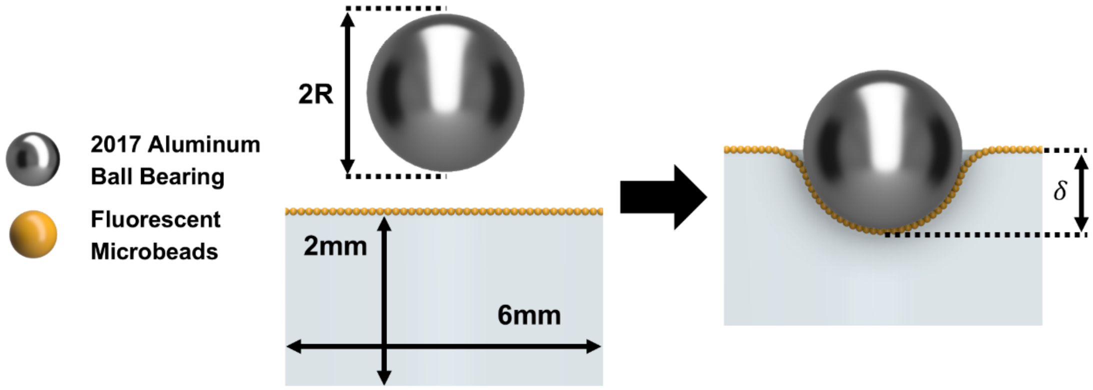


**Hertz Model**

$$E_{gel} = \frac{3(1 - v_{gel}^2)\rho_{ball}\left(\frac{4}{3}\pi R^3\right) g}{4\sqrt{R}\delta^{3/2}}$$

R = ball radius
$\rho_{ball}$ = ball density
$g$ = 9.81 m/s$^2$
$h_0$ = initial hydrogel height
$h_i$ = height after indentation
$\delta = h_0 - h_i$
$v_{gel}$ = 0.3
$E_{app}$ = Apparent Young's Modulus

**Figure S3.** Optical macroindentation setup and Hertz model derivation for hydrogel mechanical characterization. Schematic detailing the macroindentation methodology used to assess the apparent Young's modulus($E_{app}$) of the soft, adherent bioprinted constructs. Hydrogel discs (6 mm diameter, 2 mm height) were surface-coated with 1.0 µm red fluorescent microbeads. A 2017 aluminum ball bearing (1 mm diameter) was centered on the construct to induce deformation.

Confocal Z-stack imaging (using a 10× objective) was utilized to optically quantify the indentation depth ($\delta$) by measuring the difference between the undeformed hydrogel surface height ($h_0$) and the indented surface height ($h_i$).The apparent Young's modulus was subsequently calculated using the provided modified Hertz model equation, which incorporates the known volume and density ($\rho$) of the ball bearing to determine applied force, the measured indentation depth ($\delta$), and an assumed Poisson's ratio ($\nu$) of 0.3 for the hydrogel network. Representative confocal Z-projections (bottom) illustrate the optical tracking of the fluorescent microbeads (orange) to accurately resolve construct deformation.

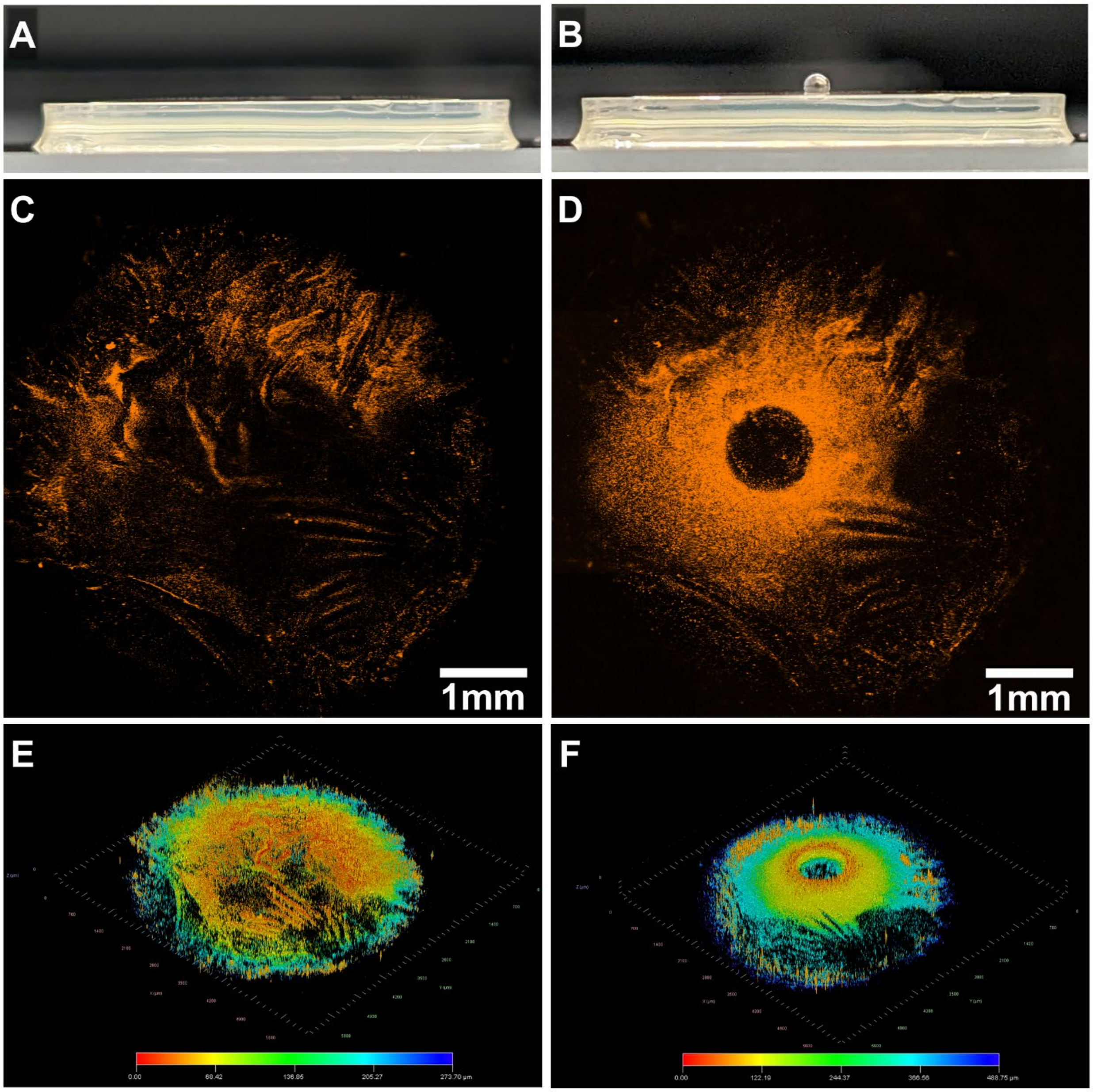


**Figure S4.** Visual and topographical validation of hydrogel deformation during macroindentation. (A, B) Representative macroscopic side-profile photographs of the bioprinted 4D hydrogel (A) in a resting state and (B) undergoing active compression with a 2017 aluminum ball bearing. (C, D) Maximal confocal Z-projections of the hydrogel surface, visualized via coated fluorescent microbeads, (C) prior to indentation and (D) under ball-bearing induced deformation (scale bars = 1 mm). (E, F) Corresponding 3D isometric surface plots generated from the confocal Z-stacks, illustrating the topographical height mapping (E) used to establish the uncompressed baseline ($h_0$) and (F) used to strictly quantify the maximum indentation depth ($h_i$) for Hertz model calculations. Color maps in E and F represent the absolute Z-depth profile from 0 μm (red/orange) to the deepest focal plane (blue).

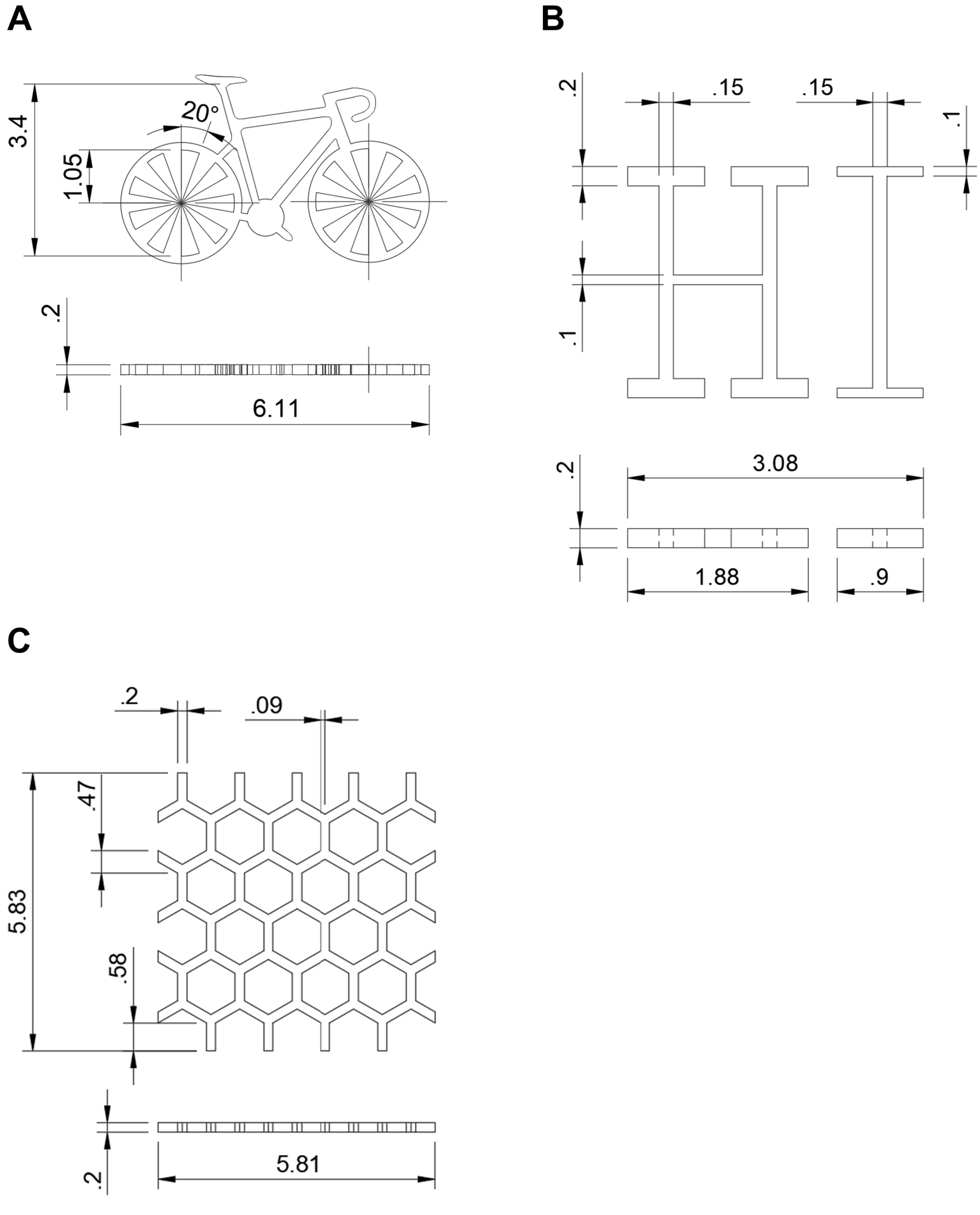


**Figure S5.** Quantitative dimensions of the computer-aided design (CAD) models utilized for print fidelity evaluation. Detailed CAD drawings corresponding to the bioprinted constructs evaluated in Figure 3. (A) The bicycle model features wheels with alternating positive and negative spaces at 20° intervals and a central spoke length of 1.05 mm, designed to assess optical scattering and determine the minimum spatial resolution of the 4D hydrogel bioink. (B) The "HI" construct details specific beam widths of 0.1 mm, 0.15 mm, and 0.2 mm to evaluate structural fidelity, post-print swelling, and mechanical stability against column collapse across varying aspect ratios. (C) The honeycomb lattice incorporates a continuous positive feature with a uniform 0.2 mm wall thickness to test macroscopic structural integrity against cupping and suction forces generated during the bioprinting process. All three models were configured for fabrication to a uniform target height of 0.2 mm (200 μm), and all annotations are presented in millimeters.

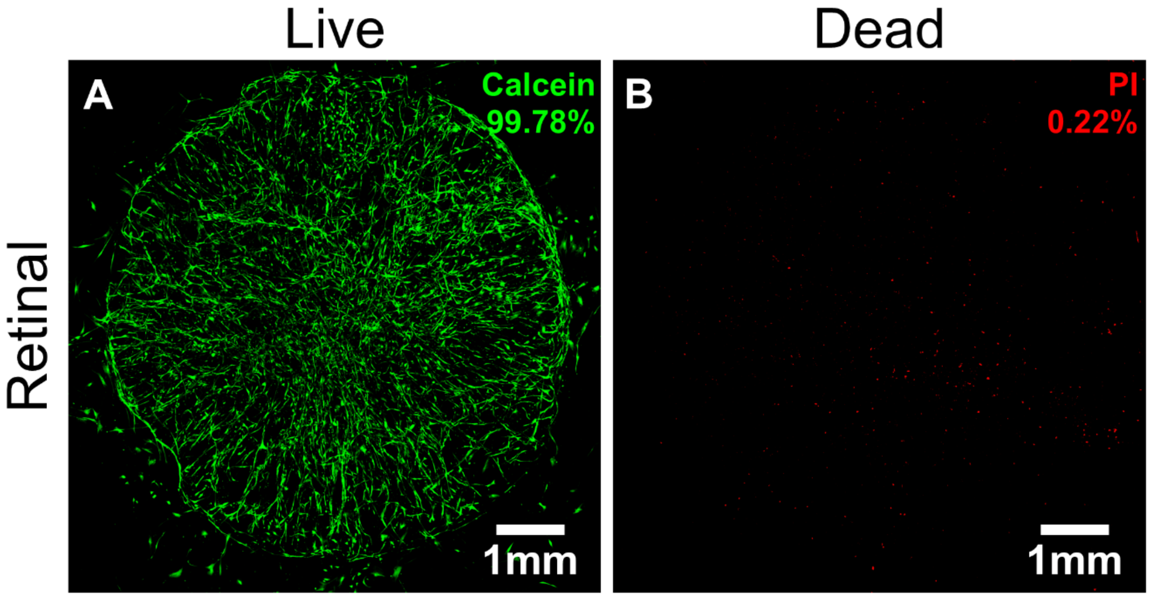


**Figure S6**. Separated fluorescence channels for live/dead viability assays of bioprinted retinal and pulmonary constructs. (A) Individual fluorescence channels for the live (Calcein-AM, green) and dead (Propidium Iodide, red) assays of the Day 7 retinal triculture model presented in Figure

4. Quantitative analysis demonstrated robust cytocompatibility with 99.78% live cells and 0.22% dead cells. All scale bars = 1 mm.

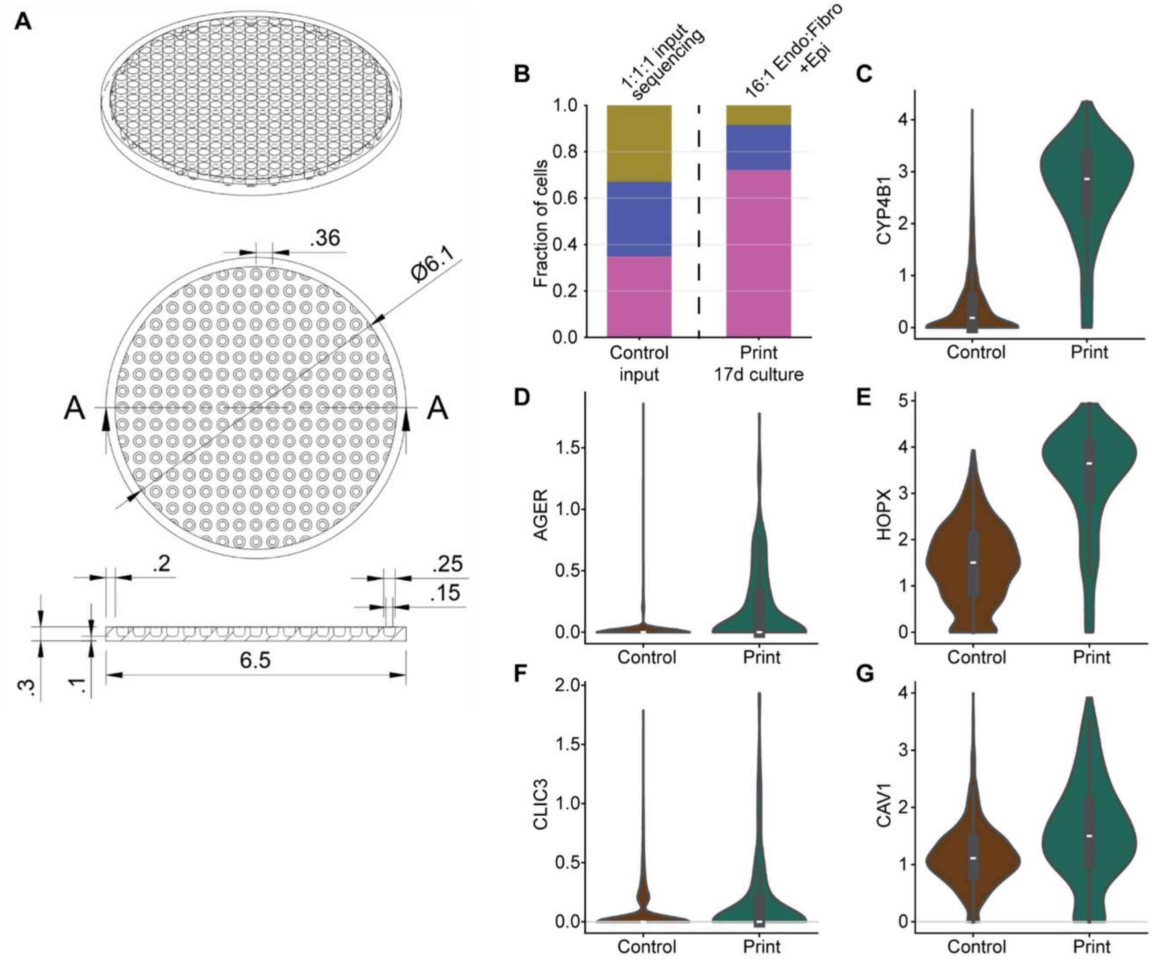


**Figure S7.** Pulmonary tissue construct characterization. A) Quantitative dimensions of the computer-aided design (CAD) lung microwell model utilized to incorporate lung organoids. Detailed CAD drawings corresponding to Figure 5. All dimensional annotations are presented in millimeters (mm). B) Stacked bar plot showing the relative proportions of endothelial, epithelial, and fibroblast compartments in the control input and bioprinted construct conditions. **(C)** Violin plot showing expression of CYP4B1, an AT1-associated epithelial marker, within the epithelial subset separated by experimental condition. AT1 epithelial signature score derived from canonical atlas-based AT1 epithelial markers, showing increased AT1-associated transcriptional enrichment within the epithelial compartment following culture in bioprinted pulmonary constructs. Violin plot showing expression of AT1 markers (D) AGER, (E) HOPX, (F) CLIC3, and (G) CAV1.

Tables

| Time (Days) | Gel Composition (w/v) | Mean Apparent Young's Modulus (Pa) | Standard Deviation | Standard Error | Tukey Grouping | | | | |
|---|---|---|---|---|---|---|---|---|---|
| 0 | 6.25% GelMA | 3705.87693 | 63.90324 | 26.08839 | A | | | | |
| 1 | 6.25% GelMA | 3633.76928 | 114.47331 | 46.73353 | A | | | | |
| 2 | 6.25% GelMA | 3609.1876 | 85.7023 | 34.98782 | A | | | | |
| 4 | 6.25% GelMA | 3638.16134 | 147.37421 | 60.16527 | A | | | | |
| 7 | 6.25% GelMA | 3642.4662 | 70.58773 | 28.81732 | A | | | | |
| 10 | 6.25% GelMA | 3676.79485 | 78.90144 | 32.21138 | A | | | | |
| 14 | 6.25% GelMA | 3721.66281 | 72.97695 | 29.79272 | A | | | | |
| 0 | 3.75%/2.5% GelMA/Gelatin | 1929.57784 | 191.07267 | 78.00509 | | B | | | |
| 1 | 3.75%/2.5% GelMA/Gelatin | 1689.07653 | 161.24979 | 65.82995 | | | C | | |
| 2 | 3.75%/2.5% GelMA/Gelatin | 1510.35037 | 121.30509 | 49.52259 | | | C | | |
| 4 | 3.75%/2.5% GelMA/Gelatin | 1193.19111 | 91.14543 | 37.20997 | | | | D | |
| 7 | 3.75%/2.5% GelMA/Gelatin | 1118.77866 | 60.48006 | 24.69088 | | | | D | E |
| 10 | 3.75%/2.5% GelMA/Gelatin | 1038.34023 | 60.09796 | 24.53489 | | | | D | E |
| 14 | 3.75%/2.5% GelMA/Gelatin | 997.33393 | 36.70237 | 14.98368 | | | | | E |

**Table S1.** Descriptive statistics and post-hoc pairwise comparisons of hydrogel apparent Young's modulus. Comprehensive statistical breakdown of the ball-bearing macroindentation data presented in Figure 2B. The table details the mean apparent Young’s modulus, standard deviation, and standard error for the 6.25% w/v GelMA control and the 4D hydrogel (3.75% w/v GelMA / 2.5% w/v gelatin) across the 14-day culture period ($n=6$ independent samples per condition). Global statistical significance was determined using a two-way analysis of variance (ANOVA). To evaluate specific differences between individual time points and formulations, Tukey’s Honestly Significant Difference (HSD) post-hoc test was applied at a 95% confidence level. Within the "Tukey Grouping" column, conditions are categorized by letter assignments to denote statistical relationships. Conditions that share at least one corresponding letter (e.g., a group labeled "DE" and a group labeled "E") exhibit no statistically significant difference in their mean stiffness. Conversely, conditions that do not share any common letters (e.g., Group "A" versus Group "B", or "C" versus "D") are statistically significantly different from one another.
